# Non-glycosylated Seipin to Cause a Motor Neuron Disease Induces ER stress and Apoptosis by Inactivating the ER Calcium Pump SERCA2b

**DOI:** 10.1101/2021.10.22.465502

**Authors:** Shunsuke Saito, Tokiro Ishikawa, Satoshi Ninagawa, Tetsuya Okada, Kazutoshi Mori

**Author notes:** To whom correspondence should be addressed: Kazutoshi Mori, Department of Biophysics, Graduate School of Science, Kyoto University, Kitashirakawa-Oiwake, Sakyo-ku, Kyoto 606-8502, Japan. Biosignal Research Center, Kobe University, 1-1, Rokkodai-cho, Nada-ku, Kobe 657-8501, Japan (Satoshi Ninagawa). Abbreviations: Endo H, endoglycosidase H; ER, endoplasmic reticulum; ERAD, ER-associated degradation; IP3R, IP3 receptor; ngSeipin, KO, knock out; non-glycosylated Seipin; PLA, proximity ligation assay; RyR, Ryanodine Receptor; TM, transmembrane; TPM, transcripts per million; UPR, unfolded protein response; Tg, thapsigargin; WT, wild-type; wtSeipin, wild-type Seipin; 4CmC, 4-chloro-m-cresol.

## Abstract

A causal relationship between endoplasmic reticulum (ER) stress and the development of neurodegenerative diseases remains controversial. Here, we focused on Seipinopathy, a dominant motor neuron disease, based on the finding that its causal gene product, Seipin, is a hairpin-like transmembrane protein in the ER. Gain-of-function mutations of Seipin produce non-glycosylated Seipin (ngSeipin), which was previously shown to induce ER stress and apoptosis at both cell and mouse levels albeit with no clarified mechanism. We found that aggregation-prone ngSeipin dominantly inactivated SERCA2b, the major calcium pump in the ER, and decreased the calcium concentration in the ER, leading to ER stress and apoptosis. This inactivation required oligomerization of ngSeipin and direct interaction of the ngSeipin C-terminus with SERCA2b, and was observed in Seipin-deficient human colorectal carcinoma-derived cells (HCT116) expressing ngSeipin at a level comparable with that in neuroblastoma cells (SH-SY5T). Our results thus provide a new direction to controversy noted above.

## INTRODUCTION

The *BSCL1* and *BSCL2* genes have been identified as causal genes of congenital generalized lipodystrophy (CGL) or Berardinelli-Seip congenital lipodystrophy syndrome (BSCL), a rare autosomal recessive disease characterized by insufficiency of adipose tissue from birth or early infancy and by severe insulin resistance. The *BSCL1* gene encodes 1-acylglycerol-3-phosphate O-acyltransferase 2 (AGPAT2), which is present in the membrane of the endoplasmic reticulum (ER) and involved in phospholipid biosynthesis (Agarwal et al., 2002; Garg et al., 1999), whereas the *BSCL2* gene encodes Seipin, a hairpin-like transmembrane protein in the ER with unknown function at that time (Magre et al., 2001). Loss of function mutations of the *BSCL2* gene appear to produce more severe symptoms than those of the *BSCL1* gene (Van Maldergem et al., 2002). Since the discovery that Seipin is involved in lipid droplet morphology in yeast (Szymanski et al., 2007), the role of Seipin in the biogenesis of lipid droplets has gained extensive attention (Bi et al., 2014; Cui et al., 2011; Sim et al., 2013; Sui et al., 2018; Tian et al., 2011; Wang et al., 2014; Wang et al., 2016; Yan et al., 2018).

To our interest, two missense mutations of the *BSCL2* gene, namely N152S and S154L of Seipin, were found to dominantly cause distal hereditary motor neuropathy (dHMN) or distal muscular atrophy, which is characterized by almost exclusively degenerated motor nerve fibers, predominantly in the distal part of limbs (Windpassinger et al., 2004). Because N^152^, V^153^ and S^154^ of Seipin match the triplet code (Asn-X-Ser/Thr; X: any amino acid except Pro) for N-glycosylation, neither N152S nor S154L Seipin are glycosylated, leading to the proposal that the production of these aggregation-prone mutants results in neurodegeneration (Windpassinger et al., 2004).

Seipin was first identified as a protein of 398 aa (Magre et al., 2001), and was later found (Lundin et al., 2006) to have two splice variants, a short form of 398 aa and long form of 462 aa (see Fig. 1A), which are translated from three Seipin mRNA isoforms of 1.6 kb, 1.8 kb, and 2.2 kb. Both forms are translatable from 1.8 kb and 2.2 kb mRNA but the long form is more abundantly produced than the short form. In contrast, only the short form is translated from 1.6 kb mRNA (Lundin et al., 2006). Because 1.8 kb mRNA is predominantly expressed in human brain (Magre et al., 2001), it is considered that human brain expresses mainly the long form (Cartwright and Goodman, 2012).

**Fig. 1.**
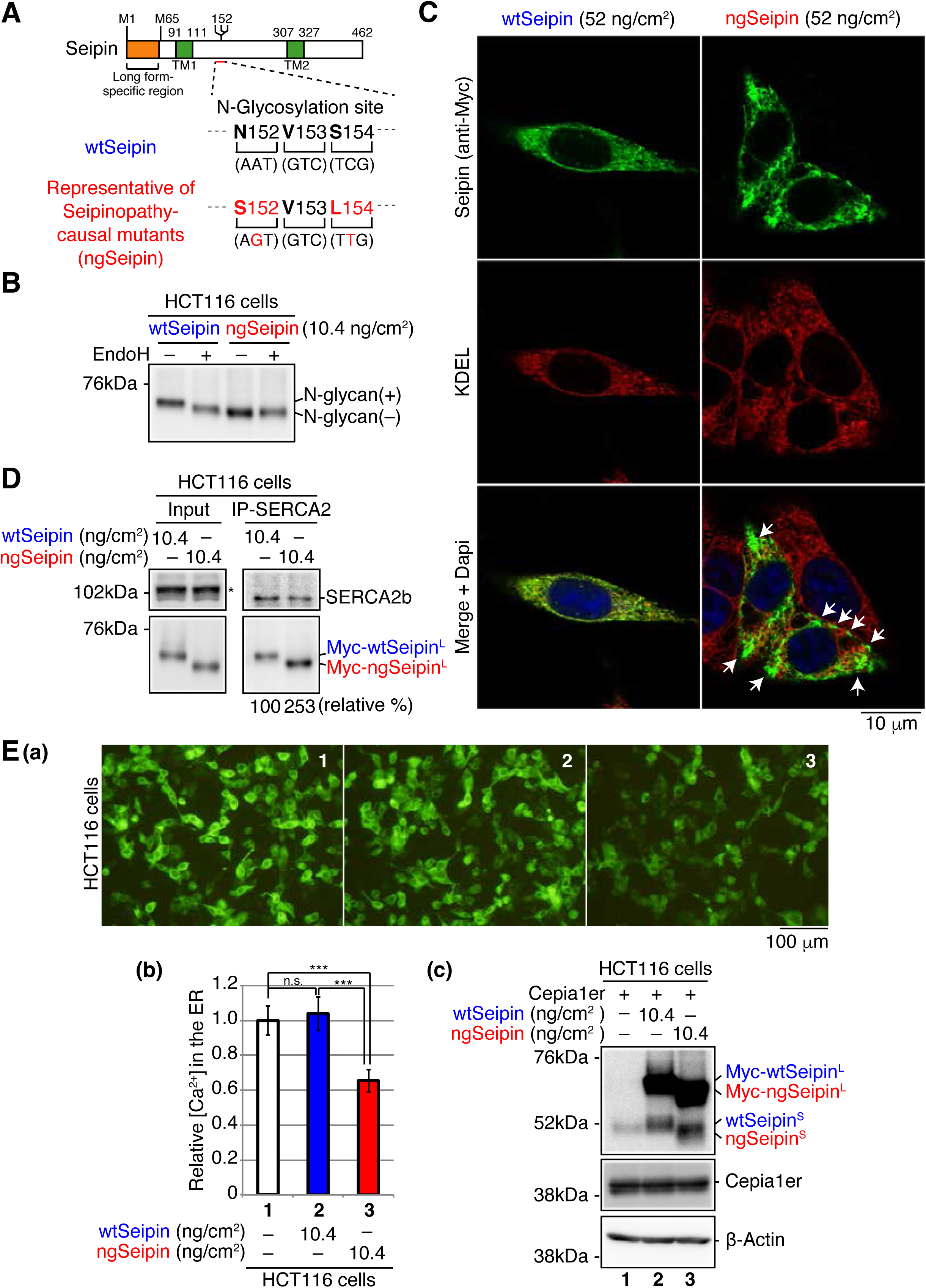
Effect of ngSeipin expression on calcium concentration in the ER of HCT116 cells. (A) Structure of human Seipin is schematically shown. The orange and green boxes denote the long form-specific region and transmembrane (TM1 and TM2) domains, respectively. The amino acid sequences of 152-154 and corresponding nucleotide sequences of wild-type Seipin (wtSeipin) and representative of Seipinopathy-causal mutants (non-glycosylated Seipin, ngSeipin) are shown below. (B) Cell lysates were prepared from HCT116 cells transfected with plasmid (10.4 ng/cm^2^) to express Myc-tagged wtSeipin or ngSeipin (both long forms), treated with (+) or without (-) EndoH, and analyzed by immunoblotting using anti-Myc antibody. (C) HCT116 cells transfected with plasmid (52 ng/cm^2^) to express Myc-tagged wtSeipin or ngSeipin were analyzed by immunofluorescence using anti-Myc and anti-KDEL antibodies, which recognized major ER chaperones. Scale bar: 10 μm. White arrowheads indicate aggregated ngSeipin. (D) Cell lysates were prepared from HCT116 cells transfected with plasmid (10.4 ng/cm^2^) to express Myc-tagged wtSeipin or ngSeipin, and subjected to immunoprecipitation using anti-SERCA2 antibody. Aliquots of cell lysates (Input) and immunoprecipitates (IP-SERCA2) were analyzed by immunoblotting using anti-SERCA2 and anti-Myc antibodies. The asterisk denotes a non-specific band. (E) HCT116 cells were transfected with plasmid (104 ng/cm^2^) to express Cepia1er together with or without plasmid (10.4 ng/cm^2^) to express Myc-tagged wtSeipin or ngSeipin. (a) Fluorescent microscopic analysis of transfected cells was conducted. Scale bar: 100 μm. (b) Fluorescence intensities of 16-17 pictures obtained from three independent experiments (5-7 pictures each) were quantified and are expressed relative to that in cells transfected with plasmid to express Cepia1er alone with error bars (standard deviation). (c) Cell lysates were prepared and analyzed by immunoblotting using anti-Seipin, anti-GFP and anti-*β*-actin antibodies.

The ER, where Seipin is located, is well known to be equipped with a quality control system for proteins. Productive folding of newly synthesized secretory and transmembrane proteins is assisted by ER-localized molecular chaperones and folding enzymes (ER chaperones hereafter). In contrast, proteins unable to gain their correct three-dimensional structures are dealt with by ER-associated degradation (ERAD), in which unfolded or misfolded proteins are recognized, delivered to the transmembrane complex termed the retrotranslocon, and retrotranslocated to the cytosol for ubiquitin-dependent proteasomal degradation.

Under a variety of physiological and pathological conditions, however, this quality control system misfunctions, resulting in the accumulation of unfolded or misfolded proteins in the ER. This ER stress is quite detrimental to the cell and may eventually cause cell death. In response, ER stress is immediately and adequately counteracted by a cellular homeostatic mechanism termed the unfolded protein response (UPR). In vertebrates, the UPR is triggered by three types of ubiquitously expressed ER stress sensor and transducers - PERK, ATF6 and IRE1 - which leads to general translational attenuation to decrease the burden on the ER; transcriptional upregulation of ER chaperons to increase productive folding capacity; and transcriptional upregulation of ERAD components to increase degradation capacity.

Daisuke Ito and colleagues showed for the first time that expression of non-glycosylated mutant Seipin by transfection in HeLa cells evokes ER stress, as evidenced by induction of the two major ER chaperones BiP and GRP94, the ERAD component Herp, and CHOP. HeLa cells expressing non-glycosylated mutant Seipin by transfection are subject to more extensive apoptosis (18%) than those expressing wild-type (WT) Seipin (6%) (Ito and Suzuki, 2007). Based on these findings, they proposed the designation of mutant Seipin-linked dominant motor neuron disease as Seipinopathy, which represents a novel ER stress-associated disease (Ito and Suzuki, 2009).

They further constructed a transgenic mice overexpressing human non-glycosylated mutant Seipin under the control of the neuron-specific murine Thy-1 promoter. They found that the levels of ER stress marker proteins BiP and PDI are elevated in brain of the transgenic mice, reproducing the symptomatic and pathological phenotypes observed in human patients with Seipinopathy (Yagi et al., 2011).

Here, we focused on the remaining and most critical question of how non-glycosylated mutant Seipin evokes ER stress.

## RESULTS

### Construction of Seipinopathy-causal mutant Seipin

We found that human HCT116 diploid cells derived from colorectal carcinoma (Roschke et al., 2002), which we use exclusively for gene knockout analysis, expressed only the short form of Seipin, designated Seipin^S^, whereas human neuroblastoma-derived SH-SY5Y cells with trisomy 7 (Yusuf et al., 2013) expressed both Seipin^S^ and the long form of Seipin, designated Seipin^L^, and that Seipin^S^ and Seipin^L^ were both sensitive to digestion with endoglycosidase H (Endo H) [Fig. 1-S1A(a)]. When normalized by level of GAPDH, which was expressed quite similarly in HCT116 and SH-SY5Y cells [Fig. 1-S1A(b)], quantitative RT-PCR showed that SH-SY5Y cells expressed Seipin mRNA 5 times more abundantly than HCT116 cells [Fig. 1-S1A(c)].

To produce the representative of Seipinopathy-causal mutants, we simultaneously mutated Asn^152^ and Ser^154^ of Seipin^L^ to Ser and Leu, respectively (Fig. 1A). When expressed in HCT116 cells by transfection, N-terminally Myc-tagged WT Seipin^L^ was sensitive to digestion with Endo H but N-terminally Myc-tagged mutant (N152S/S154L) Seipin^L^ was not, as expected (Fig. 1B); WT Seipin^L^ and the non-glycosylated mutant Seipin^L^ are hereafter designated wtSeipin^L^ and ngSeipin^L^, respectively. wtSeipin^L^ expressed in HCT116 cells by transfection showed typical ER pattern, as expected, whereas ngSeipin^L^ showed uneven distribution in the ER, suggesting that ngSeipin is prone to aggregation (Fig. 1C). It should be noted that as we carried out transfection in cell culture systems of various sizes, we express a transfection index as the amount of plasmid (ng) divided by the bottom area (cm^2^) of the well/dish, i.e. 2.0 cm^2^ for a 24-well plate, 9.6 cm^2^ for a 6-well plate, and 11.8 cm^2^ for a 3.5-cm dish, for easier comparison of results obtained from different experiments; accordingly, when 100 and 123 ng plasmid was transfected into cells in 6-well plates and 3.5 cm dishes, respectively, the transfection index was 10.4 ng/cm^2^.

### Effect of ngSeipin expression on calcium concentration in the ER

We focused on SERCA2, the major calcium pump in the ER incorporating cytosolic calcium ion into the ER, based on the previous observation that Seipin physically associates with SERCA in fly as well as with SERCA2 in HEK293 cells (Bi et al., 2014). It should be noted that the expression level of SERCA2 dominated that of SERCA1 and SERCA3 in both HCT116 and SH-SY5Y cells (Fig. 1-S1B). It is also known that three splice variants exist for SERCA2 (Gélébart et al., 2003) and that SERCA2b is ubiquitously expressed, whereas SERCA2a and SERCA2c are expressed mainly in myocardium and skeletal muscle, in which the expression level of SERCA2b is low (Dally et al., 2006). Indeed, the expression level of SERCA2b dominated that of SERCA2a and SERCA2c in both HCT116 and SH-SY5Y cells (Fig. 1-S1C). Interestingly, immunoprecipitation from HCT116 cells expressing wtSeipin^L^ or ngSeipin^L^ by transfection showed that ngSeipin^L^ bound to SERCA2b more extensively than wtSeipin^L^ (Fig. 1D).

To monitor the calcium concentration ([Ca^2+^] hereafter) in HCT116 cells, we employed the fluorescent reporters Cepia1er and GCaMP6f, whose fluorescence reflects [Ca^2+^] in the ER (Suzuki et al., 2014) and in cytosol (Chen et al., 2013), respectively. Expression of ngSeipin^L^ in HCT116 cells by transfection (10.4 ng/cm^2^) markedly decreased [Ca^2+^] in the ER compared with that of wtSeipin^L^ (Fig. 1E). This effect of ngSeipin^L^ on [Ca^2+^] in the ER was also observed in SH-SY5Y cells (Fig. 1-S1D). We noticed that small amounts (∼20%) of wtSeipin^S^ and ngSeipin^S^ were produced from transfected plasmid [Fig. 1E(c); and see Fig. 3A(c) for clearer evidence]. Because they were not detected with anti-Myc antibody (data not shown), it is likely that they were translated from the second methionine M^65^ (see Fig. 1A), given that the nucleotide sequences around M65 (ccgGccATGG) are more similar to the Kozak consensus sequence for translational initiation (gccRccATGG) than those around M1 (aggAagATGt).

### Specific and dominant inactivation of SERCA2b by ngSeipin

The ER contains two types of calcium channel, namely Ryanodine Receptor (RyR) and IP3 receptor (IP3R), which release stored calcium to cytosol upon various stimuli, for example 4-chloro-m-cresol (4CmC) for RyR (Zorzato et al., 1993) and bradykinin for IP3R (Cruzblanca et al., 1998). Quantitative RT-PCR detected expression of RyR1, IP3R1, IP3R2 and IP3R3 in HCT116 cells (Fig. 2-S1A). 4CmC, Bradykinin, or both decreased [Ca^2+^] in the ER in untransfected HCT116 cells and in HCT116 cells expressing wtSeipin^L^ by transfection (10.4 ng/cm^2^), as expected [Fig. 2-S1B(d), white and blue bars]. The lowered [Ca^2+^] in the ER of HCT116 cells expressing ngSeipin^L^ compared with those expressing wtSeipin^L^ [Fig. 2-S1B(d); compare bars 5, 11, and 17 with bars 3, 9, and 15] was further decreased upon treatment with 4CmC, bradykinin, or both [Fig. 2-S1B(d); compare bar 6 with bar 5, bar 12 with bar 11, bar 18 with bar 17], suggesting that RyR and IP3R are still active in HCT116 cells expressing ngSeipin^L^.

Thapsigargin treatment rapidly increases [Ca^2+^] in cytosol by inhibiting SERCA1/2/3 without affecting RyR and IP3R (Lytton et al., 1991). Total amount of calcium released from the ER to cytosol upon thapsigargin treatment, which was monitored using GCaMP6f, was markedly decreased in HCT116 cells expressing ngSeipin^L^ by transfection (10.4 ng/cm^2^) compared with those expressing wtSeipin^L^ (Fig. 2A), reflecting lowered [Ca^2+^] in the ER (Fig. 1E), which was monitored using Cepia1er. Treatment of HCT116 cells or HCT116 cells expressing wtSeipin^L^ by transfection (10.4 ng/cm^2^) with CDN1163, a SERCA2 activator (Cornea et al., 2013; Gruber et al., 2014), increased [Ca^2+^] in the ER, as expected, whereas treatment of HCT116 cells expressing ngSeipin^L^ by transfection with CDN1163 did not do (Fig. 2-S2A). These results suggest that ngSeipin^L^ selectively inactivates SERCA2b.

**Fig. 2.**
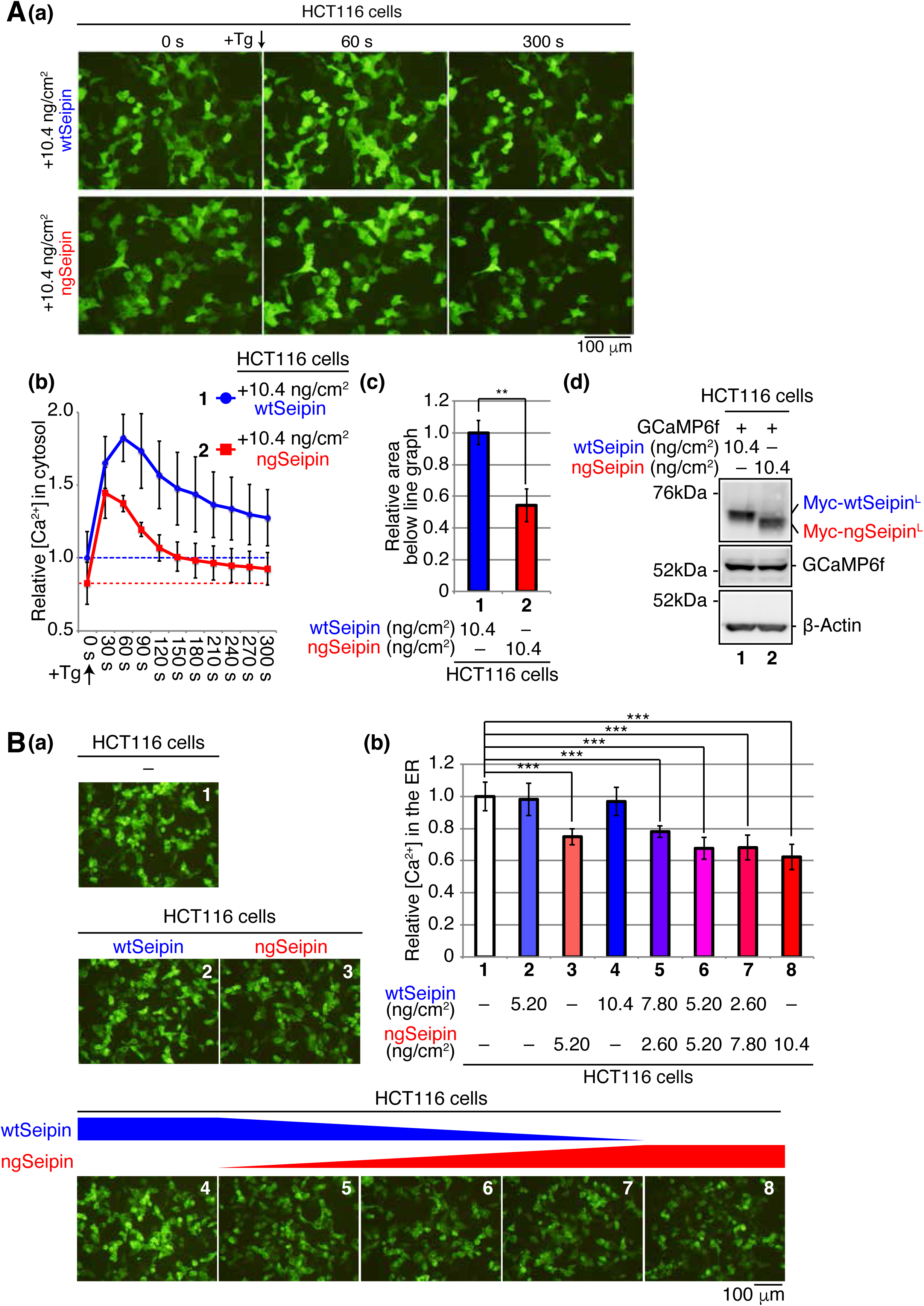
Effect of co-expression of ngSeipin and wtSeipin on calcium concentration in the ER of HCT116 cells. (A) HCT116 cells were transfected with plasmid (104 ng/cm^2^) to express GCaMP6f together with plasmid (10.4 ng/cm^2^) to express Myc-tagged wtSeipin or ngSeipin. (a) Fluorescent microscopic analysis was conducted before (0 s) and every 30 s after treatment with 1 μM thapsigargin (Tg). Only pictures of 0 s, 60 s and 300 s are shown. Scale bar: 100 μm. (b) Fluorescence intensities were quantified at each time point and are shown as a line graph with the fluorescence intensity in HCT116 cells expressing wtSeipin before Tg treatment set as 1 (n=3). (c) The area below the red line graph until the broken red line (fluorescence intensity at 0 s, +ngSeipin) in (b) was calculated and is shown relative to that below the blue line graph until the broken blue line (fluorescence intensity at 0 s, +wtSeipin). (d) Cell lysates were prepared and analyzed by immunoblotting using anti-Myc, anti-GFP and anti-*β*-actin antibodies. (B) HCT116 cells were transfected with plasmid (104 ng/cm^2^) to express Cepia1er together with or without indicated amounts of plasmid to express Myc-tagged wtSeipin or ngSeipin. (a) Fluorescent microscopic analysis was conducted. Scale bar: 100 μm. (b) Fluorescence intensities were quantified and are expressed as in Fig. 1E(b) (n=3).

**Fig. 3.**
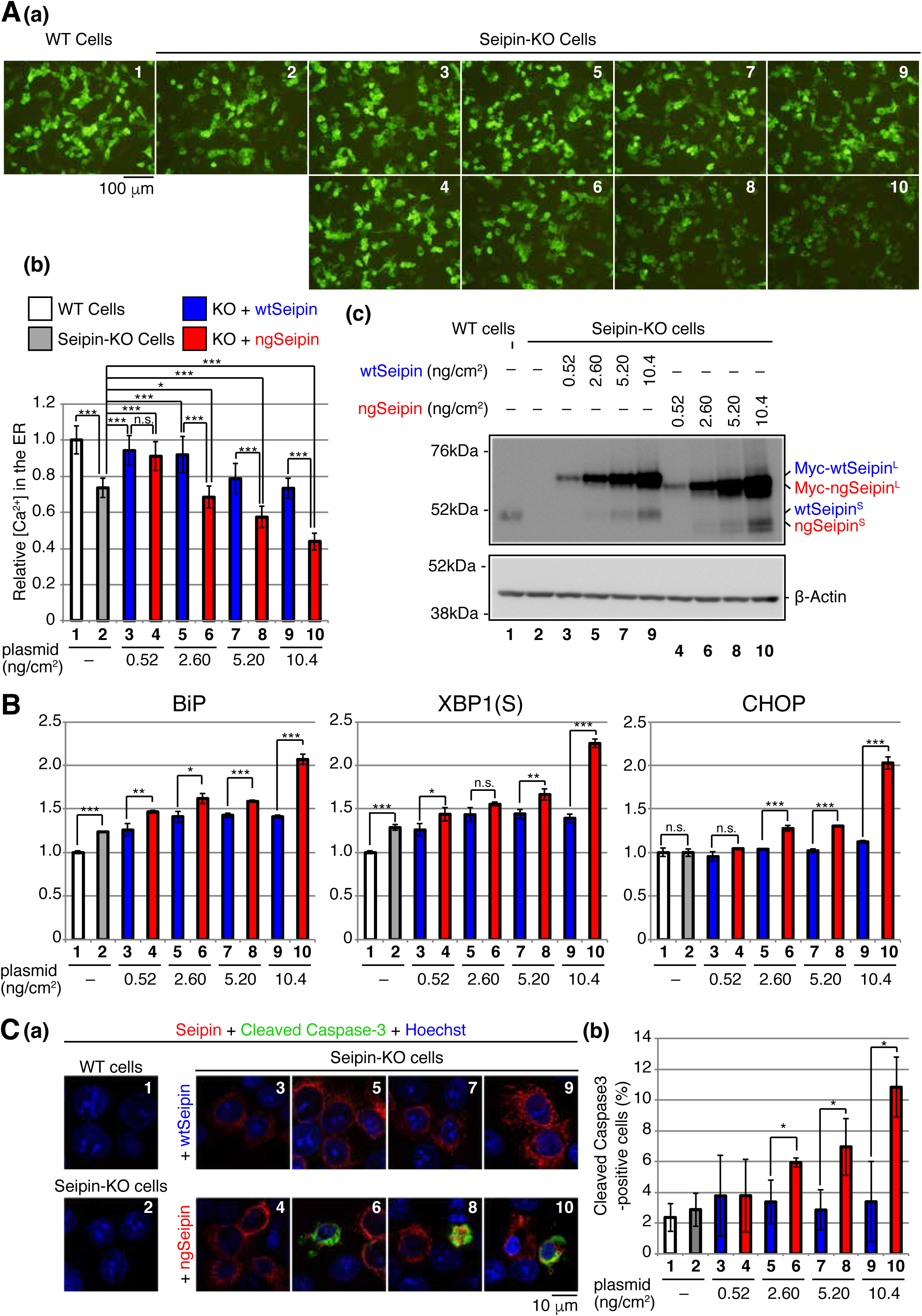
Effect of ngSeipin expression on calcium concentration in the ER, ER stress and apoptosis in Seipin-KO cells. (A) WT cells were transfected with plasmid (104 ng/cm^2^) to express Cepia1er. Seipin-KO cells were transfected with plasmid (104 ng/cm^2^) to express Cepia1er together with or without indicated amounts of plasmid to express Myc-tagged wtSeipin or ngSeipin. (a) Fluorescent microscopic analysis of WT cells and Seipin-KO cells transfected as indicated was conducted. Scale bar: 100 μm. (b) Fluorescence intensities were quantified and are expressed as in Fig. 1E(b) (n=3). (c) Cell lysates were prepared from the indicated cells and analyzed by immunoblotting using anti-Seipin and anti-*β*-actin antibodies. (B) Quantitative RT-PCR was conducted to determine the levels of endogenous BiP mRNA, spliced XBP1 [XBP1(S)] mRNA and CHOP mRNA relative to that of GAPDH mRNA in WT cells and Seipin-KO cells transfected with the indicated amounts of plasmid to express Myc-tagged wtSeipin or ngSeipin (n=3). The mean value of untransfected WT cells is set as 1. (C) (a) WT cells, Seipin-KO cells, and Seipin-KO cells transfected with the indicated amount of plasmid to express Flag-tagged wtSeipin or ngSeipin were fixed after 28 h, subjected to immunostaining using anti-Flag and anti-Cleaved Caspase-3 antibodies, and analyzed by confocal microscopy. Scale bar: 10 μm. (b) The number of Flag-tagged Seipin (red) and Cleaved Caspase-3 (green) double-positive cells was counted in 118-250 cells obtained from three independent experiments and are shown as a percentage.

Because Seipinopathy is an autosomal dominant disease, we next examined the effect of co-expression of wtSeipin^L^ and ngSeipin^L^ on [Ca^2+^] in the ER. Expression of ngSeipin^L^ by transfection at 5.20 and 10.4 ng/cm^2^ decreased [Ca^2+^] in the ER in a dose-dependent manner [Fig. 2B(b); compare bars 3 and 8 with bar 1], whereas expression of wtSeipin^L^ by transfection at 5.20 and 10.4 ng/cm^2^ did not do [Fig. 2B(b); compare bars 2 and 4 with bar 1]. Co-expression of wtSeipin^L^ in a decreasing manner by transfection at 7.80, 5.20, and 2.60 ng/cm^2^ and of ngSeipin^L^ in an increasing manner by transfection at 2.60, 5.20, and 7.80 ng/cm^2^ (transfection a total of 10.4 ng/cm^2^) decreased [Ca^2+^] in the ER [Fig. 2B(b); compare bars 5, 6 and 7 with bar 1]. Furthermore, CDN1163 treatment did not increase [Ca^2+^] in the ER significantly in HCT116 cells co-expressing wtSeipin^L^ and ngSeipin^L^ by transfection (5.20 ng/cm^2^ each), in contrast to the case of untransfected HCT116 cells (Fig. 2-S2B). Thus, ngSeipin^L^ dominantly inactivates SERCA2b and thereby dominantly decreases [Ca^2+^] in the ER.

### Construction and characterization of Seipin-knockout cells

To examine the effect of endogenous Seipin on SERCA2b, we knocked out (KO) the *Seipin* gene in HCT116 cells using CRISPR/Cas9-mediated cleavage of the *Seipin* locus at two sites (Fig. 3-S1A). The deletion of almost the entire *Seipin* gene was confirmed by genomic PCR (Fig. 3-S1B), and the absence of *Seipin* mRNA and Seipin protein was confirmed by RT-PCR (Fig. 3-S1C) and immunoblotting [Fig. 3A(c); compare lane 2 with lane 1], respectively.

[Ca^2+^] in the ER was decreased in Seipin-KO cells by 20∼30% compared with WT cells [Fig. 3A(b); compare bar 2 with bar 1], consistent with an approximately 30% decrease in SERCA activity in Seipin mutant fly compared with WT fly and with Bi et al.’s proposal that Seipin helps maintenance of calcium homeostasis in the ER by binding to SERCA (Bi et al., 2014). In addition, oleic acid (400 μM)-induced lipid drop formation was suppressed in Seipin-KO cells compared with WT cells [Fig. 3-S1D(c); compare dot plot 2 with dot plot 1], as reported previously (Chung et al., 2019; Salo et al., 2016). Expression of both wtSeipin^L^ and ngSeipin^L^ in Seipin-KO cells at a low level by transfection at 0.52 ng/cm^2^, which was comparable with the level of endogenous Seipin^S^ [Fig. 3A(c); compare lanes 3 and 4 with lane 1], restored [Ca^2+^] in the ER [Fig. 3A(b); compare bars 3 and 4 with bar 1], and rescued the phenotype of Seipin-KO cells in terms of oleic acid-induced lipid drop formation [Fig. 3-S1D(c); compare dot plots 4 and 5 with dot plot 1].

In contrast, a higher expression of ngSeipin^L^ in Seipin-KO cells by transfection at ≥ 2.60 ng/cm^2^ decreased [Ca^2+^] in the ER in dose-dependent manner more robustly than that of wtSeipin^L^ [Fig. 3A(b); compare bars 6, 8 and 10 with bars 5, 7 and 9]. Higher expression of ngSeipin^L^ by transfection at ≥ 2.60 ng/cm^2^ induced ER stress more extensively than that of wtSeipin^L^ in Seipin-KO cells, as evidenced by increased levels of BiP mRNA (a target of the ATF6 pathway), XBP1(S) mRNA (a target of the IRE1 pathway) and CHOP mRNA (a target of the PERK pathway) (Fig. 3B; compare bars 6, 8, and 10 with bars 5, 7 and 9). Higher expression of ngSeipin^L^ by transfection at ≥ 2.60 ng/cm^2^ induced apoptosis more extensively than that of wtSeipin^L^ in Seipin-KO cells, as shown by increased detection of Cleaved Caspase-3 by immunofluorescence (Fig. 3C(b); compare bars 6, 8, and 10 with bars 5, 7 and 9). Of note, clear-cut difference in the effect on [Ca^2+^] in the ER and apoptosis between transfection of ngSeipin at 0.52 ng/cm^2^ (not significant) and ≥ 2.60 ng/cm^2^ (significant) [Fig. 3A(b) and Fig. 3C(b)] is most well reflected in induction of CHOP (Fig. 3B), suggesting the importance of the PERK pathway in this ER stress-induced apoptosis. These results suggest that ngSeipin^L^ expression-mediated decrease in [Ca^2+^] in the ER would be a key for the development of Seipinopathy.

### Effect of oligomerization of ngSeipin on inactivation of SERCA2b

To elucidate the mechanism by which ngSeipin^L^ inactivates SERCA2b, we examined the effect of oligomerization of Seipin, because human Seipin exists as a wheel-like undecamer (Fig. 4A) (Yan et al., 2018). Interestingly, fly Seipin consisting of 370 aa is not glycosylated and exists as a wheel-shaped dodecamer (Sui et al., 2018). Because N-glycan wedges the interface of two protomers in the case of human Seipin (Fig. 4A, right panel), we hypothesized that non-glycosylated human Seipin expressed at a higher level becomes unable to maintain the undecamer structure, leading to aggregation.

**Fig. 4.**
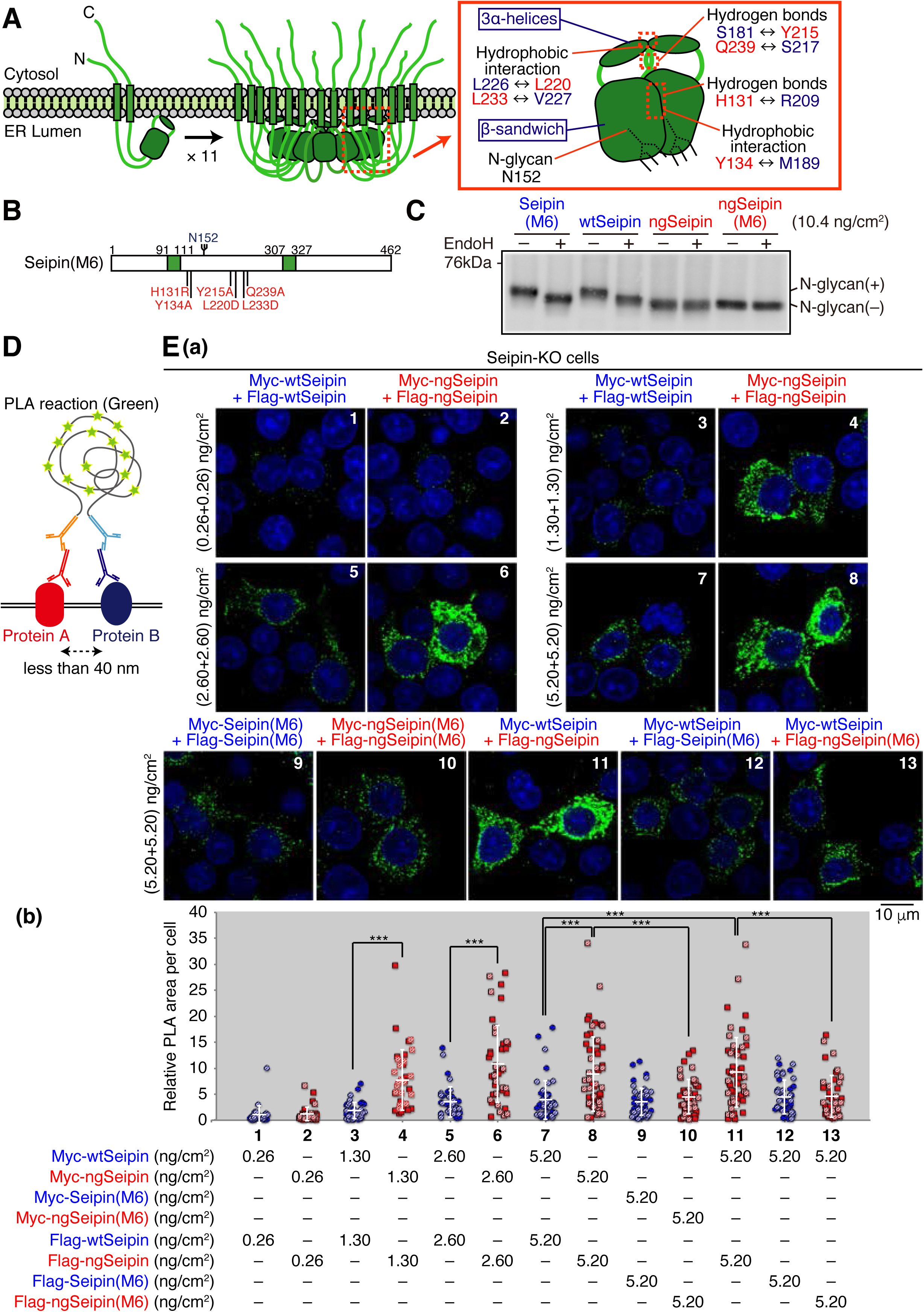
Luminal region-mediated aggregation of ngSeipin. (A) Structures of human Seipin monomer and undecamer are schematically shown on the left. Structure of one Seipin protomer and its neighboring protomer is schematically shown on the right with the positions of the *β*-sandwich domain, three *α*-helices, hydrophobic interactions, hydrogen bonds, and N-glycan. (B) Structure of Seipin(M6) is schematically shown, in which the six amino acids critical for oligomerization of Seipin are mutated and highlighted in red. (C) Cell lysates were prepared from WT cells transfected with plasmid (10.4 ng/cm^2^) to express Myc-tagged wtSeipin, Seipin(M6), ngSeipin or ngSeipin(M6), treated with (+) or without (-) EndoH, and analyzed by immunoblotting using anti-Myc antibody. (D) The principle of PLA is diagrammatically presented. PLA detects the proximal interaction (less than 40 nm) of two proteins in the cell using two different antibodies. (E) (a) PLA was conducted in Seipin-KO cells transfected with the indicated amounts of plasmid to simultaneously express Myc-tagged and Flag-tagged proteins with various combinations as indicated, and analyzed by confocal microscopy. Scale bar: 10 μm. (b) PLA signals were quantified in 35-57 cells obtained from two independent experiments (filled and striped dots denote the data of experiment 1 and 2, respectively) and are shown as signals (summation of PLA-positive area) per cell relative to those observed in cells transfected simultaneously with plasmid to express Myc-tagged wtSeipin (0.26 ng/cm^2^) and plasmid to express Flag-tagged wtSeipin (0.26 ng/cm^2^).

Structural analysis revealed that the luminal region of each Seipin monomer consists of 8 *β*-strands, termed the *β*-sandwich domain, and 3 *α*-helices, and that an ER membrane-anchored core-ring is formed when the 3 *α*-helices of each of 11 monomers is gathered via multiple hydrophobic interactions between one protomer and its neighboring protomer, including those between L226 and L220 and between L233 and V227. The core ring is surrounded by 11 *β*-sandwich domains which are tightly associated via hydrogen bonds between one protomer and its neighboring protomer, including those between S181 and Y215, between Q239 and S217, and between H131 and R209, and via hydrophobic interactions between one protomer and its neighboring protomer, including those between Y134 and M189 (Fig. 4A, right panel) (Yan et al., 2018).

We therefore simultaneously mutated the 6 amino acids in wtSeipin^L^ and ngSeipin^L^ (H131R, Y134A, Y215A, L220D, L233D, and Q239A, Fig. 4B), as previously reported to prevent oligomerization (Yan et al., 2018). Resulting Seipin^L^(M6) and ngSeipin^L^(M6) were still sensitive and insensitive, respectively, to digestion with EndoH, as expected (Fig. 4C). Of note, Seipin^L^(M6) and ngSeipin^L^(M6) tended to lose the ability to rescue the phenotype of Seipin-KO cells in terms of oleic acid-induced lipid drop formation [Fig. 3-S1D(c); compare dot blots 6 and 7 with dot blot 1], suggesting the importance of Seipin oligomerization in this phenotype.

To confirm the aggregation propensity of ngSeipin, we employed a proximity ligation assay (PLA), in which a PCR-mediated signal is produced when the distance between two proteins is less than 40 nm (Fig. 4D), and which was used to detect aggregates of *α*-synuclein, a causal protein of familial Parkinson’s disease (Roberts et al., 2015). Results showed the production of a markedly strong signal when Myc-tagged ngSeipin^L^ and Flag-tagged ngSeipin^L^ were co-expressed by transfection each at ≥ 1.30 ng/cm^2^ (total at ≥ 2.60 ng/cm^2^), compared with co-expression of Myc-tagged wtSeipin^L^ and Flag-tagged wtSeipin^L^ [Fig. 4E(b); compare dot plot 4 with dot plot 3, dot plot 6 with dot plot 5, and dot plot 8 with dot plot 7], as we expected. Importantly, this signal was diminished when Myc-tagged ngSeipin^L^(M6) and Flag-tagged ngSeipin^L^(M6) were co-expressed by transfection each at 5.20 ng/cm^2^ (total at 10.4 ng/cm^2^) [Fig. 4E(b); compare dot plot 10 with dot plot 8], supporting our hypothesis of non-glycosylated Seipin oligomer (undecamer)-dependent aggregation when expressed at a higher level. Accordingly, although immunoprecipitation showed association of ngSeipin^L^(M6) with SERCA2b (Fig. 5A), ngSeipin^L^(M6) introduced by transfection even at 10.4 ng/cm^2^ (highest transfection level in this report) did not decrease [Ca^2+^] in the ER (Fig. 5B), did not induce ER stress (Fig. 5C), and did not induce apoptosis (Fig. 5D), in marked contrast to ngSeipin^L^. These results suggest that SERCA2b is incorporated into aggregates of oligomerized ngSeipin^L^ to be inactivated. Of note, Flag-tagged ngSeipin^L^ tended to form aggregates with Myc-tagged wtSeipin more extensively than Flag-tagged ngSeipin^L^(M6) when analyzed by PLA [Fig. 4E(b); compare dot plot 11 with dot blot 13], indicative of its dominant nature.

**Fig. 5.**
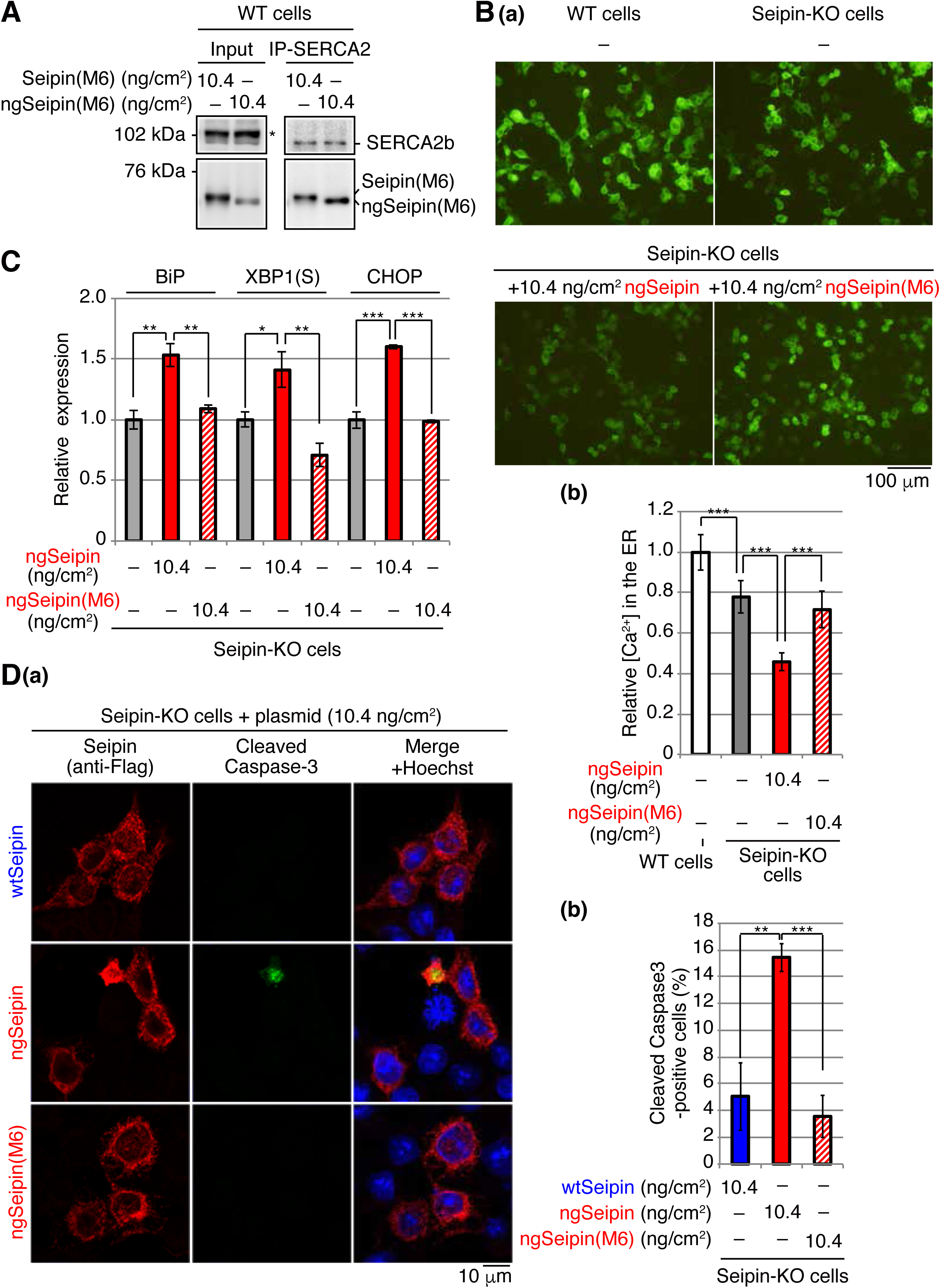
Effect of ngSeipin oligomerization on calcium concentration in the ER, ER stress and apoptosis in Seipin-KO cells. (A) Cell lysates were prepared from HCT116 cells transfected with plasmid (10.4 ng/cm^2^) to express Myc-tagged Seipin(M6) or ngSeipin(M6), subjected to immunoprecipitation using anti-SERCA2 antibody, and analyzed as in Fig. 1D. (B) WT cells were transfected with plasmid (104 ng/cm^2^) to express Cepia1er. Seipin-KO cells were transfected with plasmid (104 ng/cm^2^) to express Cepia1er together with or without plasmid (10.4 ng/cm^2^) to express Myc-tagged ngSeipin or ngSeipin(M6). (a) Fluorescent microscopic analysis was conducted. Scale bar: 100 μm. (b) Fluorescence intensities were quantified and are expressed as in Fig. 1E(b) (n=3). (C) Quantitative RT-PCR was conducted in Seipin-KO cells transfected with or without plasmid (10.4 ng/cm^2^) to express Myc-tagged ngSeipin or ngSeipin(M6) (n=3), as in Fig. 3B. (D) (a) (b) Seipin-KO cells transfected with plasmid (10.4 ng/cm^2^) to express Flag-tagged wtSeipin, ngSeipin or ngSeipin(M6) were fixed after 28 h, subjected to immunostaining, and analyzed as in Fig. 3C using 108-118 cells (n=3). Scale bar: 10 μm.

### Requirement of both the luminal and C-terminal regions of ngSeipin for inactivation of SERCA2b

To determine which region(s) of ngSeipin^L^ is required for inactivation of SERCA2b, we constructed a series of deletion mutants in wtSeipin^L^ and ngSeipin^L^, namely *Δ*N lacking the cytosolic N-terminal region, *Δ*LD lacking the luminal region, and *Δ*C lacking the cytosolic C-terminal region (Fig. 6A). We also constructed two swap mutants of wtSeipin^L^ and ngSeipin^L^, in which the first and second transmembrane (TM) domains of Seipin^L^ were replaced by the fourth and first TM domain of glucose 6-phosphatase, respectively (Fig. 6A), in reference to the previous swapping experiments (Bi et al., 2014) and with further consideration of the topology of these TM domains (Fig. 6-S1A). All constructs produced a band of the expected size in transfected WT cells (Fig. 6B and Fig. 6-S1B, Input).

**Fig. 6.**
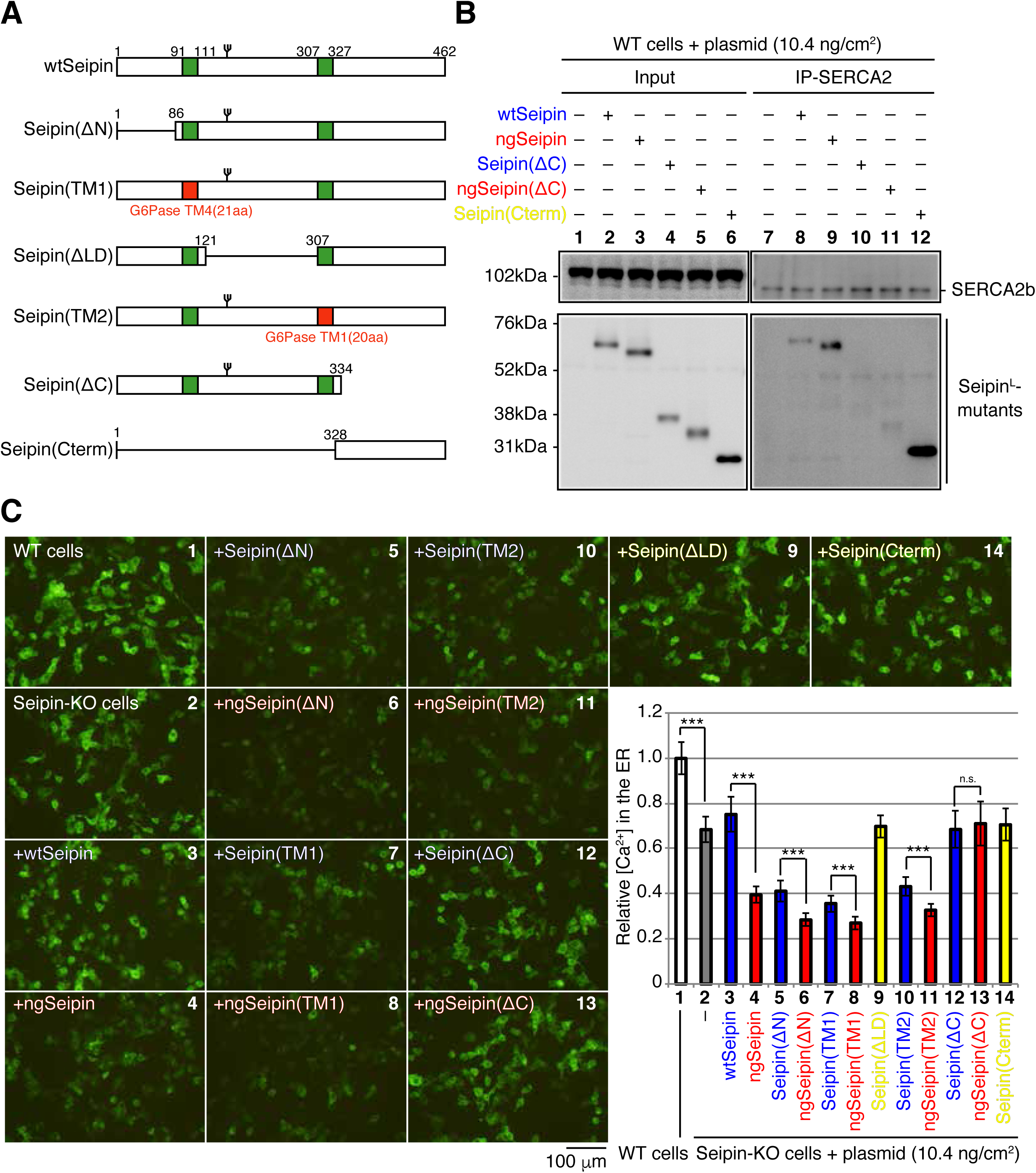
Effect of various deletions or replacements of ngSeipin on interaction with SERCA2 and calcium concentration in the ER of Seipin-KO cells. (A) Structures of wtSeipin and various mutant Seipin are schematically shown. The red boxes denote the transmembrane (TM1 and TM4) domains of G6Pase to swap the transmembrane domains of Seipin. (B) Cell lysates were prepared from WT cells transfected with plasmid (10.4 ng/cm^2^) to express Myc-tagged wtSeipin or various mutant Seipin, subjected to immunoprecipitation using anti-SERCA2 antibody, and analyzed as in Fig. 1D. (C) WT cells were transfected with plasmid (104 ng/cm^2^) to express Cepia1er. Seipin-KO cells were transfected with plasmid (104 ng/cm^2^) to express Cepia1er together with or without plasmid (10.4 ng/cm^2^) to express Myc-tagged wtSeipin wtSeipin or various mutant Seipin. (a) Fluorescent microscopic analysis of WT or Seipin-KO cells transfected as indicated was conducted. Scale bar: 100 μm. (b) Fluorescence intensities were quantified and are expressed as in Fig. 1E(b) (n=3).

Immunoprecipitation using anti-SERCA2 antibody revealed that only Seipin^L^(*Δ*C) and ngSeipin^L^(*Δ*C) were hardly co-immunoprecipitated with SERCA2b (Fig. 6B, lanes 10 and 11, and Fig. 6-S1B, lanes 23 and 24). We thus constructed a mutant which expresses only the C-terminal region of Seipin^L^ and found that this Seipin(Cterm) was efficiently co-immunoprecipitated with SERCA2b (Fig. 6B, lane 12). These findings indicate that the cytosolic C-terminal region of Seipin^L^ is necessary and sufficient for interaction with SERCA2b.

In the case of *Δ*N (Fig. 6C; compare bar 6 with bar 5), TM1 (compare bar 8 with bar 7) and TM2 (compare bar 11 with bar 10) mutants, ngSeipin^L^-based mutants decreased [Ca^2+^] in the ER more effectively than wtSeipin^L^-based mutants, similarly to the case of ngSeipin^L^ versus wtSeipin^L^ (compare bar 4 with bar 3), when expressed in Seipin-KO cells by transfection at 10.4 ng/cm^2^. In contrast, ngSeipin^L^(*Δ*C) lost the ability to decrease [Ca^2+^] in the ER (Fig. 6C; compare bar 13 with bar 12), indicating that direct interaction with SERCA2b is critical. Of note, Seipin^L^(*Δ*LD) and Seipin(Cterm) did not decrease [Ca^2+^] in the ER (Fig. 6C, yellow bars 9 and 14), although they were co-immunoprecipitated with SERCA2 (Fig. 6-S1B, lane 20 and Fig. 6B, lane 12). Given that the luminal region contains the 6 amino acids critical for oligomerization of Seipin (Fig. 4B), both the luminal and C-terminal regions of ngSeipin^L^ are required for inactivation of SERCA2b.

Interestingly, ngSeipin^L^(*Δ*C) expressed by transfection at total 10.4 ng/cm^2^ (highest transfection level in this report) existed in closer proximity with each other than Seipin^L^(*Δ*C), as shown by PLA (Fig. 7A). Nonetheless, ngSeipin^L^(*Δ*C) did not decrease [Ca^2+^] in the ER (Fig. 6C, bar 13), did not induce ER stress (Fig. 7B), and did not induce apoptosis (Fig. 7C). We concluded that the oligomerization-mediated aggregation property and direct interaction with SERCA2b are prerequisites for ngSeipin to inactivate SERCA2b and thereby induce ER stress and subsequent apoptosis (Fig. 7D).

**Fig. 7.**
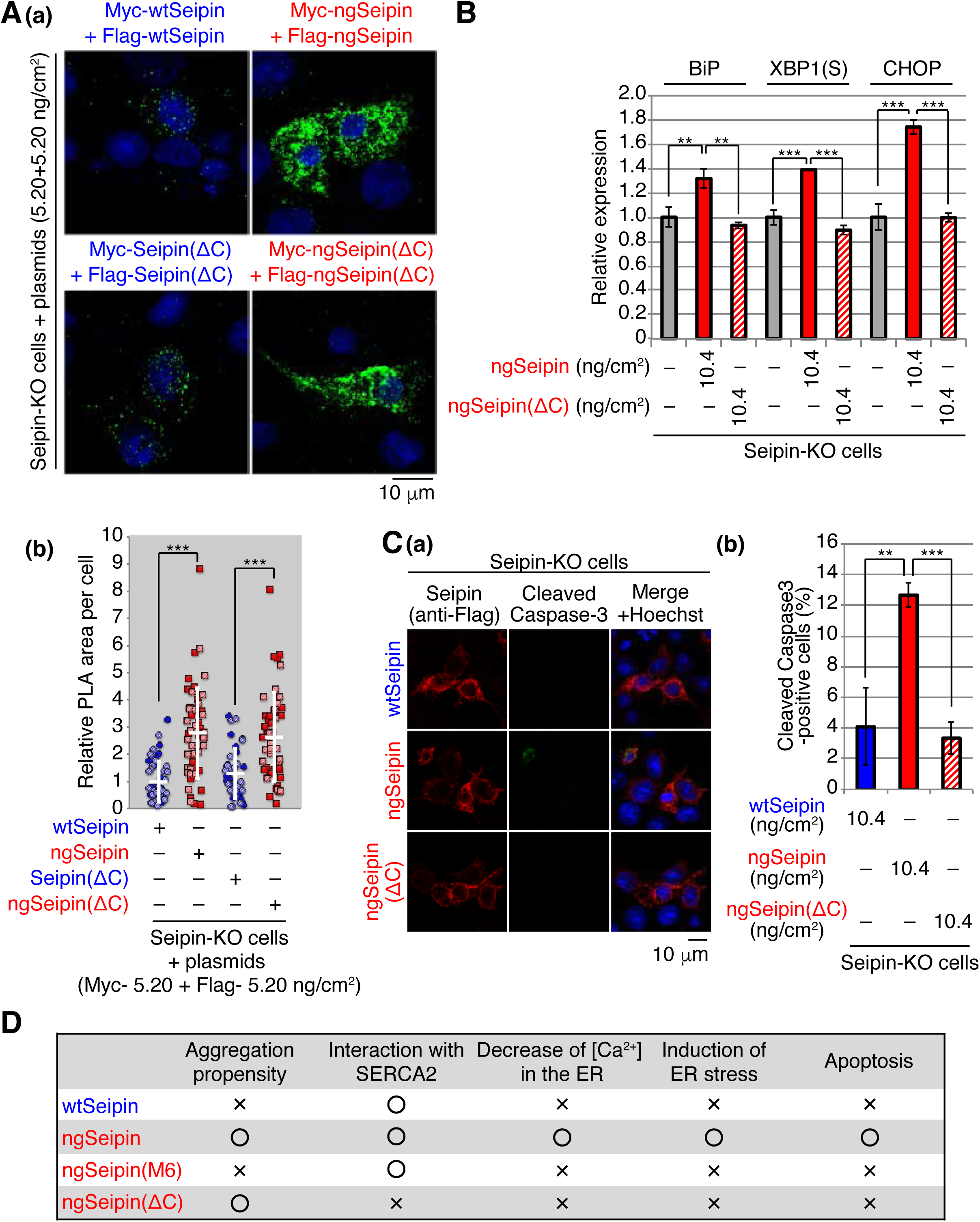
Effect of C-terminal deletion of ngSeipin on calcium concentration in the ER, ER stress and apoptosis in Seipin-KO cells. (A) (a) (b) Seipin-KO cells transfected with plasmids (5.20 ng/cm^2^ each) to simultaneously express Myc-tagged and Flag-tagged wtSeipin or various mutant Seipin as indicated were subjected to PLA and analyzed as in Fig. 4E using 40-48 cells (n=2). Scale bar: 10 μm. (B) Quantitative RT-PCR was conducted in Seipin-KO cells transfected with or without plasmid (10.4 ng/cm^2^) to express Myc-tagged ngSeipin, or ngSeipin(*Δ*C) (n=3), as in Fig. 3B. (C) (a) (b) Seipin-KO cells transfected with plasmid (10.4 ng/cm^2^) to express Flag-tagged wtSeipin, ngSeipin or ngSeipin(*Δ*C) were fixed after 28 h, subjected to immunostaining, and analyzed as in Fig. 3C using117-127 cells (n=3). Scale bar: 10 μm. (D) Phenotypes of wtSeipin, ngSeipin, ngSeipin(M6) and ngSeipin(*Δ*C) are summarized.

### Increase in the level of SERCA2b reverses the effect of ngSeipin

We finally examined whether the increase in the level of SERCA2b compensates for the decrease in [Ca^2+^] in the ER caused by expression of ngSeipin^L^. Overexpression of SERCA2b (WT) but not inactive mutant SERCA2b (Q108H) (Miyauchi et al., 2006) by transfection at 20.8 ng/cm^2^ significantly increased [Ca^2+^] in the ER of WT cells expressing ngSeipin^L^ by transfection at 10.4 ng/cm^2^ (Fig. 8A). Lowered [Ca^2+^] in the ER of Seipin-KO cells as well as of Seipin-KO cells expressing wtSeipin^L^ by transfection at 10.4 ng/cm^2^, compared with WT cells, was significantly increased by the introduction of SERCA2b in a dose-dependent manner, whereas further lowering of [Ca^2+^] in the ER of Seipin-KO cells expressing ngSeipin^L^ by transfection at 10.4 ng/cm^2^ was significantly and slightly increased only by transfection at 20.8 ng/cm^2^ of SERCA2b (Fig. 8B), indicating the severe inactivation of SERCA2b by ngSeipin^L^.

**Fig. 8.**
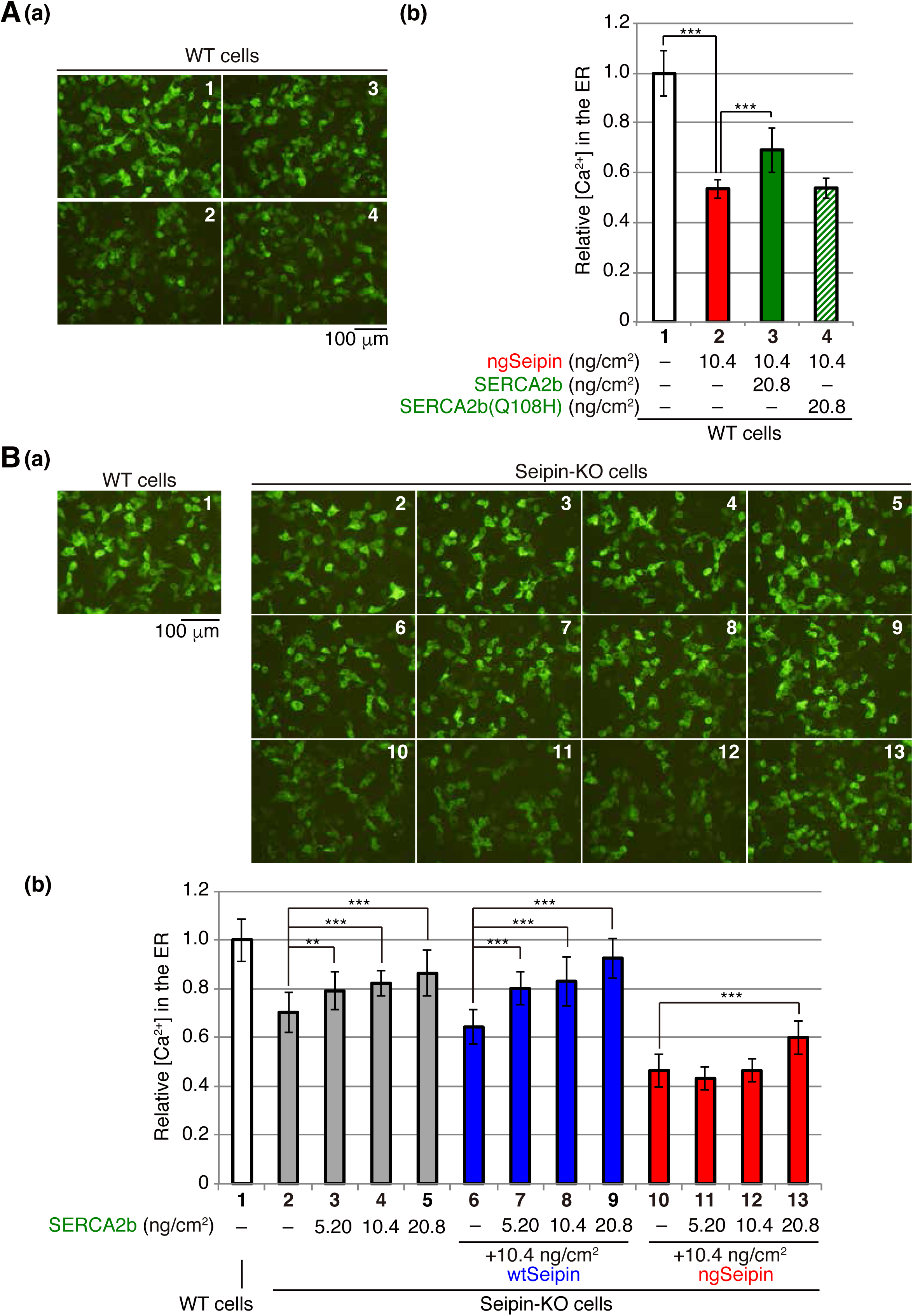
Effect of introduction of SERCA2 on calcium concentration in the ER of ngSeipin-expressing cells. (A) WT cells were transfected with plasmid (104 ng/cm^2^) to express Cepia1er together with or without the indicated amounts of plasmid to express Myc-tagged ngSeipin or Myc-tagged SERCA2b (WT or Q108H). (a) Fluorescent microscopic analysis was conducted. Scale bar: 100 μm. (b) Fluorescence intensities were quantified and are expressed as in Fig. 1E(b) (n=3). (B) WT cells were transfected with plasmid (104 ng/cm^2^) to express Cepia1er. Seipin-KO cells were transfected with plasmid (104 ng/cm^2^) to express Cepia1er together with or without the indicated amounts of plasmid to express Myc-tagged SERCA2b in addition to plasmid (10.4 ng/cm^2^) to express Myc-tagged wtSeipin or ngSeipin. (a) Fluorescent microscopic analysis was conducted. Scale bar: 100 μm. (b) Fluorescence intensities were quantified and are expressed as in Fig. 1E(b) (n=3).

Introduction of SERCA2b (transfection at 20.8 ng/cm^2^) did not affect aggregation propensity between Myc-tagged ngSeipin^L^ and Flag-tagged ngSeipin^L^ (Fig. 9A). Critically, however, introduction of SERCA2b (transfection at 20.8 ng/cm^2^) significantly mitigated the ER stress induced in Seipin-KO cells expressing ngSeipin^L^ by transfection at 10.4 ng/cm^2^ (Fig. 9B). Accordingly, the growth rate of WT cells slowed by expression of ngSeipin^L^ by transfection at 10.4 ng/cm^2^ was rescued by introduction of SERCA2b (transfection at 20.8 ng/cm^2^) (Fig. 9C). The percentage of apoptotic cells markedly increased by expression of ngSeipin by transfection at 10.4 ng/cm^2^ was greatly reduced by introduction of SERCA2b by transfection at 20.8 ng/cm^2^ (Fig. 9D). We concluded that ngSeipin induces ER stress and apoptosis through the inactivation of SERCA2b.

**Fig. 9.**
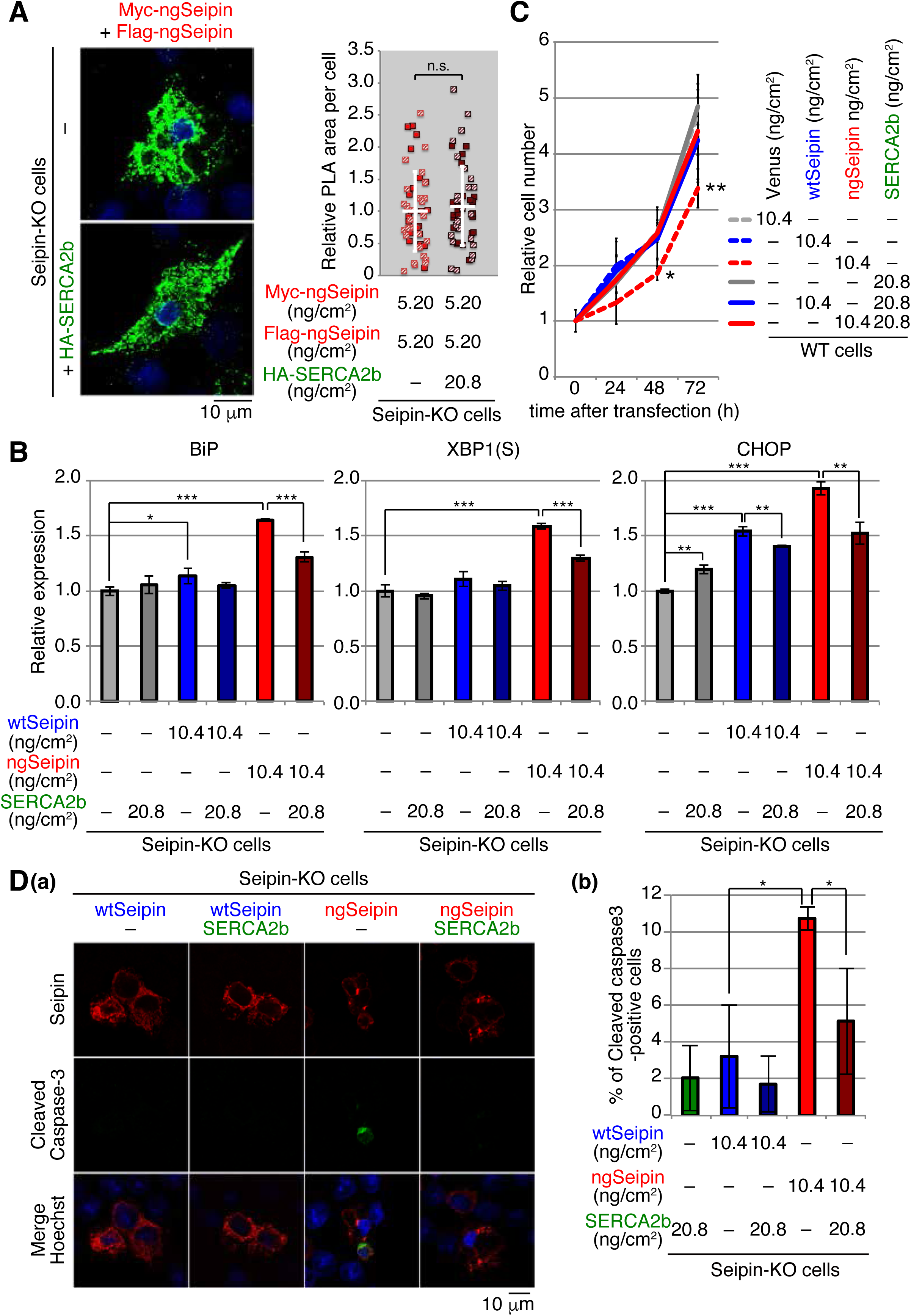
Effect of introduction of SERCA2 on ER stress, growth and apoptosis in ngSeipin-expressing cells. (A) Seipin-KO cells transfected with plasmids (5.20 ng/cm^2^ each) to simultaneously express Myc-tagged and Flag-tagged ngSeipin together with or without plasmid (20.8 ng/cm^2^) to express HA-tagged SERCA2b were subjected to PLA and analyzed as in Fig. 4E using 49-50 cells (n=2). Scale bar: 10 μm. (B) Quantitative RT-PCR was conducted in Seipin-KO cells transfected with or without plasmid (10.4 ng/cm^2^) to express Myc-tagged wtSeipin or ngSeipin together with or without plasmid (20.8 ng/cm^2^) to express Myc-tagged SERCA2b (n=3), as in Fig. 3B. (C) Growth rate of WT cells transfected with plasmid (10.4 ng/cm^2^) to express Venus, Myc-tagged wtSeipin or ngSeipin together with or without plasmid (20.8 ng/cm^2^) to express Myc-tagged SERCA2b was determined by counting cell number every 24 h (n=4). Cell number at the time of transfection is set as 1. (D) (a) (b) Seipin-KO cells transfected with plasmid (10.4 ng/cm^2^) to express Flag-tagged wtSeipin or ngSeipin together with or without plasmid (20.8 ng/cm^2^) to express Myc-tagged SERCA2b were fixed after 28 h, subjected to immunostaining, and analyzed as in Fig. 3C using 97-125 cells (n=3). Scale bar: 10 μm.

## DISCUSSION

Seipin is conserved from yeast to humans, and yeast and fly orthologues are termed Sei1 (Fld1) and dSeipin, respectively. Interestingly, Sei1 and dSeipin do not have potential N-glycosylation sites, and dSeipin was shown to function as a dodecamer of non-glycosylated monomer (Sui et al., 2018). In contrast, Seipin orthologues in vertebrates have gained one N-glycosylation site (Asn^152^Val^153^Ser^154^) in the luminal region, and one of the two N-acetylglucosamines proximal to Asn^152^ (Glc_3_Man_9_GlcNAc_2_-Asn) was shown to interact with Arg^199^ and Gly^200^ in the same molecule (Yan et al., 2018). This presence of N-glycan at the interface of each vertebrate Seipin protomer to the next vertebrate Seipin protomer in the same direction is likely to prevent dodecamer formation and instead induces undecamer formation (Fig. 4A). In this sense, non-glycosylated vertebrate Seipin is speculated to function as a dodecamer. Indeed, ngSeipin expressed at an endogenous protein level in Seipin-KO cells [transfection at 0.52 ng/cm^2^, Fig. 3A(c); compare lane 4 with lane 1] rescued [Ca^2+^] in the ER [Fig. 3A(b); compare bar 4 with bars 1 and 2] and oleic acid-induced lipid drop formation [Fig. 3-S1D(c); compare dot blot 5 with dot blots 1 and 2], both of which were mitigated in Seipin-KO cells compared with WT cells [Fig. 3A(b); compare bar 2 with bar 1, and Fig. 3-S1D(c); compare dot blot 2 with dot blot 1], suggesting that the putative dodecamer of non-glycosylated human Seipin is functional. It was previously shown that not only human wtSeipin but also ngSeipin rescued lipid droplet-associated defects in Sei1 (Fld1)-deficient yeast cells (Fei et al., 2008). These results suggest that non-glycosylation of human Seipin itself is not detrimental to the cell.

However, the presence of ngSeipin in Seipin-KO cells at a higher level (transfection at ≥ 2.60 ng/cm^2^) causes serious problems, namely a more profound decrease in [Ca^2+^] in the ER than the presence of the equivalent amount of wtSeipin [Fig. 3A(b); compare bars 6, 8 and 10 with bars 5, 7 and 9]. It should be noted that an excess level of ngSeipin decreased [Ca^2+^] in the ER of SH-SY5Y cells similarly to the case of HCT116 cells (Fig. 1-S1D), and that ngSeipin decreased [Ca^2+^] in the ER in a dominant manner over wtSeipin (Fig. 2B), consistent with its disease-causing phenotype. The decrease in [Ca^2+^] in the ER in turn induced ER stress and activation of the UPR, leading to apoptosis (Fig. 3B and 3C; compare bars 6, 8, and 10 with bars 5, 7 and 9), as ER-localized molecular chaperones require Ca^2+^ for their function; it was indeed shown that Ca^2+^ depletion destabilizes BiP-substrate complexes (Preissler et al., 2020). The decrease in [Ca^2+^] in the ER also likely detrimentally affects intracellular signaling and synaptic transmission in the nervous system.

Importantly, the 5-fold difference between 0.52 ng/cm^2^ [no indication of aggregation, Fig. 4E(b); compare dot blot 2 with dot blot 1] and 2.60 ng/cm^2^ [clear indication of aggregation, Fig. 4E(b); compare dot blot 4 with dot blot 3] is comparable with the difference in the level of Seipin mRNA between HCT116 and SH-SY5Y cells [Fig. 1-S1A(c)]. Indeed, searching the RNA sequencing database “Expression Atlas: Gene expression across species and biological conditions” (https://www.ebi.ac.uk/gxa/home) showed that Seipin mRNA is highly expressed in the nervous system compared with other tissues except for testis, whereas SERCA2 mRNA is relatively ubiquitously expressed, in humans (Fig. 10) (Consortium, 2015; Papatheodorou et al., 2020). Therefore, ngSeipin would likely easily exhibit toxic gain of function in the nervous system, although its selectivity of toxicity towards motor neurons remains to be determined.

**Fig. 10.**
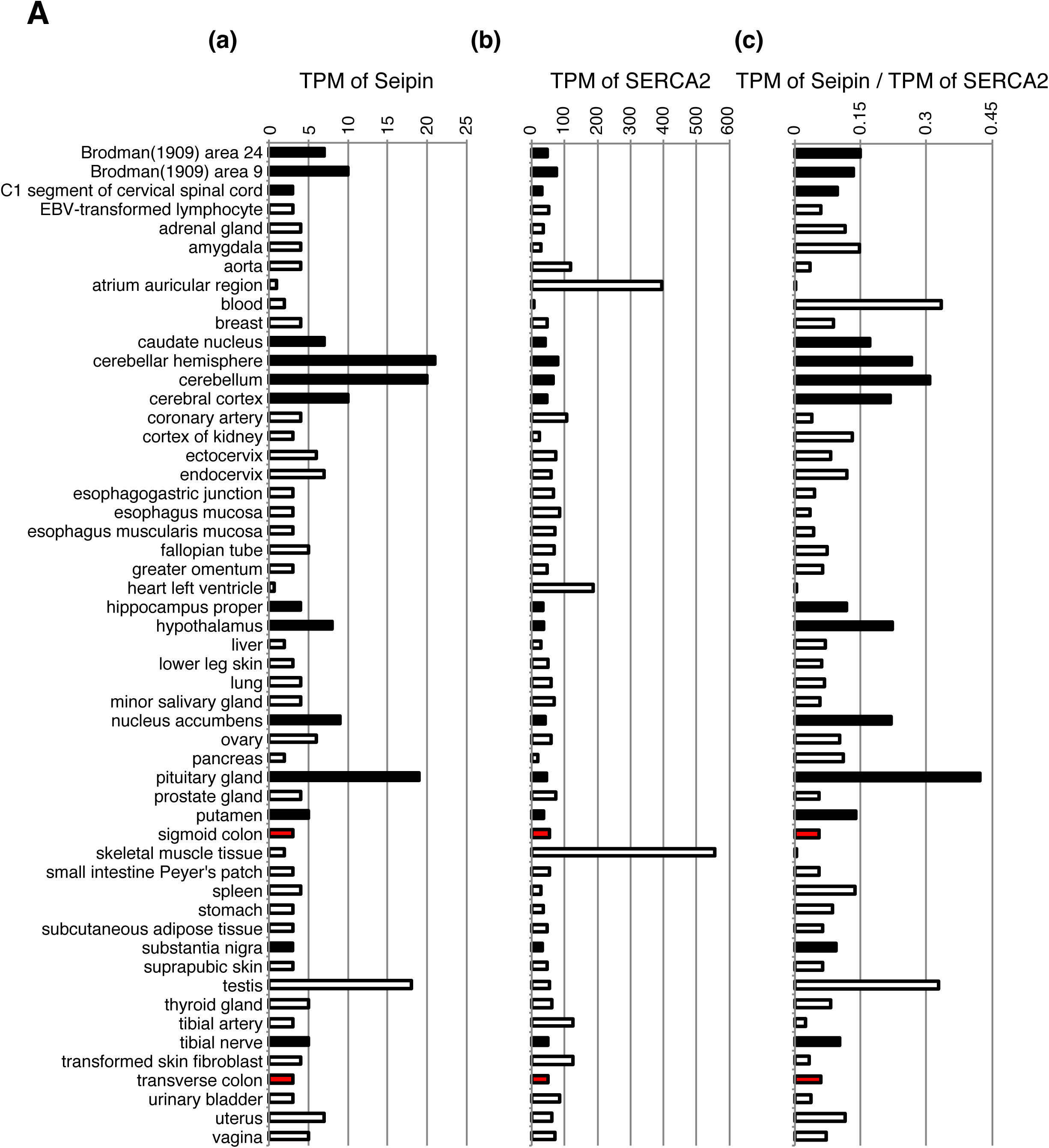
Expression levels of Seipin and SERCA2b in different human tissues. Transcripts per million (TPM) of Seipin, and TPM of SERCA2, and relative Seipin-TPM/SERCA2-TPM in different human tissues are shown. TPM data were acquired from Expression Atlas: Gene expression across species and biological conditions (https://www.ebi.ac.uk/gxa/home). Black and red bars denote data of the nervous system and of colon, respectively.

Furthermore, we unraveled the underlying molecular mechanism: ngSeipin expressed at a higher level decreases [Ca^2+^] in the ER by inactivating SERCA2b, for which two independent phenomena are prerequisite, namely oligomerization-dependent aggregation and C-terminal region-dependent direct association with SERCA2b (Fig. 7D). Thus, the prevention of oligomerization by mutating the 6 amino acids in the luminal region required for oligomerization [ngSeipin(M6), Fig. 4B] abolished the ngSeipin-mediated decrease in [Ca^2+^] in the ER (Fig. 5B), induction of ER stress (Fig. 5C), and induction of apoptosis (Fig. 5D). On the other hand, deletion of the C-terminal region of Seipin [ngSeipin(*Δ*C), Fig. 6A] abolished ngSeipin-mediated decrease in [Ca^2+^] in the ER (Fig. 6C), induction of ER stress (Fig. 7B), and induction of apoptosis (Fig. 7C), even though this ngSeipin(*Δ*C) exhibited a propensity of aggregation (Fig. 7A), indicating that the aggregation of ngSeipin is not sufficient to induce ER stress.

Importantly, the increase in the level of SERCA2b mitigated ngSeipin-mediated induction of ER stress (Fig. 9B) and subsequent apoptosis (Fig. 9D) even though ngSeipin was still severely aggregated (Fig. 9A). This raises the intriguing possibility that a potent SERCA2b activator could be used as a therapeutic for Seipinopathy and other ER stress-associated neurodegenerative diseases. In this connection, it was reported that high levels of ER stress markers were observed in motor neurons derived from patients with amyotrophic lateral sclerosis, a severe motor neuron disease, carrying the A4V mutation in superoxide dismutase 1 (Kiskinis et al., 2014; Wainger et al., 2014); and that oral administration of sodium phenylbutyrate and taurursodiol, chemical chaperones which mitigate ER stress, significantly slowed functional decline in patients with amyotrophic lateral sclerosis (Paganoni et al., 2021; Paganoni et al., 2020). These findings highlight the increasing importance of ER stress in understanding the development of certain neurodegenerative diseases

## MATERIALS AND METHODS

### Key Resources Table

| Reagent type or resources | Designation | Source or reference | Identifier | Additional information |
| --- | --- | --- | --- | --- |
| Cell line<br>(*Homo Sapiens*) | colorectal carcinoma | ATCC | HCT116 | The cell line has been tested negative for mycoplasma. |
| Cell line<br>(*Homo Sapiens*) | neuroblastoma | ATCC | SH-SY5Y | The cell line has been tested negative for mycoplasma. |
| Recombinant DNA reagent | pCMV-Myc | CLONTECH |  |  |
| Recombinant DNA reagent | pCMV G-Cepia1er | Addgene |  |  |
| Recombinant DNA reagent | pGP-CMV-GCaMP6f | Addgene |  |  |
| Recombinant DNA reagent | px330-U6-Chimeric_BB-CBh-hSpCas9 | Addgene |  |  |
| Antibody | Anti-BSCL2(Seipin) (rabbit monoclonal) | Cell Signaling | Cat#: 23846 | WB (1:1000) |
| Antibody | anti-SERCA2 (mouse monoclonal) | Santa Cruz | Cat#: sc-376235 | WB (1:500)<br>IP (5µl) |
| Antibody | anti-Myc-tag mAb-HRP-Direct (mouse monoclonal) | MBL | Code: M047-7 | WB (1:1000) |
| Antibody | anti-GFP (rabbit polyclonal) | MBL | Code: 598 | WB (1:1000) |
| Antibody | anti-β-actin (mouse monoclonal) | Wako | Cat#: 017-24573 | WB (1:2000) |
| Antibody | anti-GAPDH (rabbit polyclonal) | R&D SYSTEMS | Cat#: 2275-PC-100 | WB (1:2000) |
| Antibody | anti-KDEL (Mouse monoclonal) | MBL | Code: M181-3 | Immunostaining (1:200) |
| Antibody | anti-Myc-tag (rabbit polyclonal) | MBL | Code: 562 | PLA (1:250)<br>Immunostaining (1:100) |
| Antibody | Anti-Flag M2 (mouse monoclonal) | Sigma | Cat#: F3165 | PLA (1:250)<br>Immunostaining (1:500) |
| Antibody | Anti-Myc tag (mouse monoclonal) | Wako | Cat#: 017-21871 | Immunostaining (1:100) |
| Antibody | Anti-Cleaved Caspase-3 (rabbit polyclonal) | Cell Signaling | Cat#: 9661 | Immunostaining (1:800) |
| Antibody | Alexa488-conjugated anti-rabbit secondary antibody | Invitrogen | Cat#: A-11008 | Immunostaining (1:1000) |
| Antibody | Alexa568-conjugated anti-mouse secondary antibody | Invitrogen | Cat#: A-11004 | Immunostaining (1:1000) |
| Antibody | Alexa488-conjugated anti-mouse secondary antibody | Invitrogen | Cat#: A-11001 | Immunostaining (1:1000) |
| Antibody | Alexa594-conjugated anti-rabbit secondary antibody | Invitrogen | Cat#: A-11072 | Immunostaining (1:1000) |

### Statistics

Statistical analysis was conducted using Student’s t-test, with probability expressed as *p<0.05, **p<0.01 and ***p<0.001 for all figures. n.s. denotes not significant.

### Construction of plasmids

Recombinant DNA techniques were performed according to standard procedures (Sambrook et al., 1989) and the integrity of all constructed plasmids was confirmed by extensive sequencing analyses. pCMV-Myc (CLONTECH) was used to express a protein (Seipin, SERCA2b or Venus) tagged with Myc at the N-terminus. Site-directed mutagenesis was carried out using DpnI. Plasmids to express Flag-tagged and HA-tagged proteins were created by changing the Myc-coding sequence in pCMV-Myc expression plasmids to intended tag-coding sequence using inverse PCR and NE Builder HiFi Assembly (New England BioLabs), respectively.

### Cell culture and transfection

HCT116 cells (ATCC CCL-247) were cultured in Dulbecco’s modified Eagle’s medium (glucose 4.5 g/liter) supplemented with 10% fetal bovine serum, 2 mM glutamine, and antibiotics (100 U/ml penicillin and 100 μg/ml streptomycin) at 37°C in a humidified 5% CO2/95% air atmosphere. SH-SY5Y cells (ATCC CRL-2266) were cultured in 1:1 mixture of Eagle’s Minimum Essential Medium (glucose 1.0 g/liter) and F12 medium (glucose 1.0 g/liter) supplemented with 10% fetal bovine serum, 2 mM glutamine, and antibiotics (100 U/ml penicillin and 100 μg/ml streptomycin) at 37°C in a humidified 5% CO_2_/95% air atmosphere. Transfection was performed using polyethylenimine Max (Polyscience) according to the manufacturer’s instructions.

### Immunoblotting

HCT116 cells cultured in a 6-well plate or 3.5-cm dish were harvested with a rubber policeman and collected by centrifugation at 5,000 rpm for 2 min. Cell pellets were lysed in 200 µl of SDS sample buffer (50 mM Tris/HCl, pH 6.8, containing 100 mM dithiothreitol, 2% SDS and 10% glycerol) containing protease inhibitor cocktail (Nacalai Tesque) and 10 μM MG132. Immunoblotting analysis was carried out according to the standard procedure (Sambrook et al., 1989). Chemiluminescence obtained using Western Blotting Luminol Reagent (Santa Cruz Biotechnology) was detected using an LAS-3000mini LuminoImage analyzer (Fuji Film). EndoH was obtained from Calbiochem.

### Immunoprecipitation

Immunoprecipitation was performed using anti-SERCA2 antibody and protein A-coupled Sepharose beads (GE Healthcare). Beads were washed twice with high salt buffer (50 mM Tris/Cl, pH 8.0, containing 1% NP-40 and 150 mM NaCl), washed with PBS, and boiled for 5 min in SDS sample buffer.

### Immunofluorescence imaging

For immunofluorescence imaging, cells grown on coverslips were transiently transfected with plasmid to express Myc- or Flag-tagged protein. After 28 h cells were fixed, permeabilized with methanol at -30°C for 6.5 min, incubated at 37°C for 2 h with rabbit anti-Myc polyclonal antibody and mouse anti-KDEL monoclonal antibody, or with rabbit anti-Cleaved Caspase-3 polyclonal antibody and mouse anti-Flag monoclonal antibody, and then with Alexa488-conjugated anti-rabbit secondary antibody and Alexa568-conjugated anti-mouse secondary antibody, or Alexa488-conjugated anti-mouse secondary antibody and Alexa594-conjugated anti-rabbit secondary antibody at 37°C for 1 h. Coverslips were mounted with VECTASHIELD Mounting Medium (Vector Laboratories), Prolong Gold, or Prolong Grass Antifade Mountants (both from ThermoFisher Scientific) containing 5 µg/ml Hoechst 33342 or 50 µg/ml Dapi. Images were acquired using DM IRE2 and confocal software (both from Leica), or LSM 880 with Airyscan and Zen/Zen2.6 acquisition software (both from Carl Zeiss).

### Detection of [Ca^2+^] in the ER or cytosol

Imaging of G-Cepia1er and GCaMP6f signals were performed under a fluorescence stereomicroscope (Olympus IX-71-22TFL/PH) with acquisition software (DP Controller 1.2.1.108). Image analysis was performed using ImageJ (https://imagej.nih.gov/ij/). Bradykinin, 4-chloro-m-cresol and CDN1163 were obtained from Abcam, Tokyo Chemical Industry, and Sigma-Aldrich, respectively.

### CRISPR/Cas9 method to generate KO cell lines

PuroR fragment amplified by PCR from DT-A-pA-loxP-PGK-Puro-pA-loxP (Ninagawa et al., 2014) was inserted into the PciI site of px330-U6-Chimeric_BB-CBh-hSpCas9 (Addgene) to create px330-PuroR. The DNA oligonucleotides 5’-CACCGCTCTCACTTTCCGCCATTAG-3’ and 5’-AAACCTAATGGCGGAAAGTGAGAGC-3’, and 5’-CACCGGGGAGTGGGAAAGCTTGCTA-3’ and 5’-AAACTAGCAAGCTTTCCCACTCCCC-3’ to express gRNA for cleavage of exon 2 and the 3’UTR-non-coding region, respectively, of the *BSCL2/Seipin* gene were annealed and inserted into the BbsI site of px330-PuroR separately. HCT116 cells were co-transfected with these two plasmids using polyethylenimine Max and screened for puromycin (0.5 µg/ml) resistance.

### Genomic PCR

Non-homologous end joining in HCT116 cells was confirmed by genomic PCR using KOD-FX Neo (TOYOBO) and a pair of primers, all del Fw and all del Rv, and inside Fw and inside Rv.

### RT-PCR

Total RNA prepared from cultured cells (∼5 × 10^6^ cells) by the acid guanidinium/phenol/chloroform method using ISOGEN (Nippon Gene) was converted to cDNA using Moloney murine leukemia virus reverse transcription (Invitrogen) and oligo-dT primers. A part of the cDNA sequence of Seipin and GAPDH was amplified using PrimeSTAR GXL DNA polymerase (Takara Bio) and pairs of primers, namely Seipin cDNA Fw and Seipin cDNA Rv, and GAPDH cDNA Fw and GAPDH cDNA Rv, respectively.

### Quantitative RT-PCR

Total RNA extracted as above was subjected to quantitative RT-PCR analysis using the SYBR Green method (Applied Biosystems) and a pair of primers, namely qBiP Fw and qBiP Rv for *BiP* mRNA, qXBP1 Fw and qXBP1 Rv for spliced *XBP1* mRNA, qCHOP Fw and qCHOP Rv for *CHOP* mRNA, qGAPDH Fw and qGAPDH Rv for *GAPDH* mRNA, qRyR1 Fw and qRyR1 Rv for *RyR1* mRNA, qIP3R1 Fw and qIP3R1 Rv for *IP3R1* mRNA, qIP3R2 Fw and qIP3R2 Rv for *IP3R2* mRNA, qIP3R3 Fw and qIP3R3 Rv for *IP3R3* mRNA, qSERCA1 Fw and qSERCA1 Rv for *SERCA1* mRNA, qSERCA2 Fw and qSERCA2 Rv for *SERCA2* mRNA, qSERCA3 Fw and qSERCA3 Rv for *SERCA3* mRNA, qSERCA2abc Fw and qSERCA2a Rv for *SERCA2a* mRNA, qSERCA2abc Fw and qSERCA2b Rv for *SERCA2b* mRNA, and qSERCA2abc Fw and qSERCA2c Rv for *SERCA2c* mRNA. 200, 2,000 and 20,000 molecules of plasmid carrying Seipin, GAPDH, SERCA1, SERCA2, SERCA3, SERCA2a, SERCA2b, SERCA2c, RyR1, IP3R1, IP3R2 or IP3R3 were used as standards.

### PLA

For PLA assay, cells grown on a glass-bottom dish were transiently transfected with plasmids to express Myc-tagged and Flag-tagged proteins simultaneously. After 24 h cells were fixed and permeabilized with methanol at -30°C for 6.5 min. PLA was performed using Duolink in situ Starter Set GREEN (Sigma-Aldrich) according to manufacturer’s instructions using rabbit anti-Myc polyclonal antibody and mouse anti-Flag monoclonal antibody.

### Determination of cell growth rate

Cells transfected with various plasmids were treated with trypsin, and equal amounts of detached cells were plated to 4 dishes each. Cell numbers were counted 0, 24, 48, and 72 h later.

### Induction and imaging of lipid droplets

Cells grown on coverslips were transiently transfected with various plasmids and incubated 24 h later with medium containing 400 µM of fatty acid-free BSA (FUJIFILM)-coupled oleic acid (Nacalai Tesque). After 24 h, cells were fixed with 3.7% paraformaldehyde at -30°C for 6.5 min, and incubated with PBS containing 5.0×10^-5^% Nile Red (FUJIFILM) at 37°C for 30 min. Coverslips were mounted with Prolong Glass Antifade Mountants (ThermoFisher Scientific) containing 5 µg/ml Hoechst33342. Images were acquired using LSM 880 with Airyscan and Zen/Zen2.6 acquisition software (both from Carl Zeiss). Image analysis was performed using ImageJ.

### Supplementary File 1

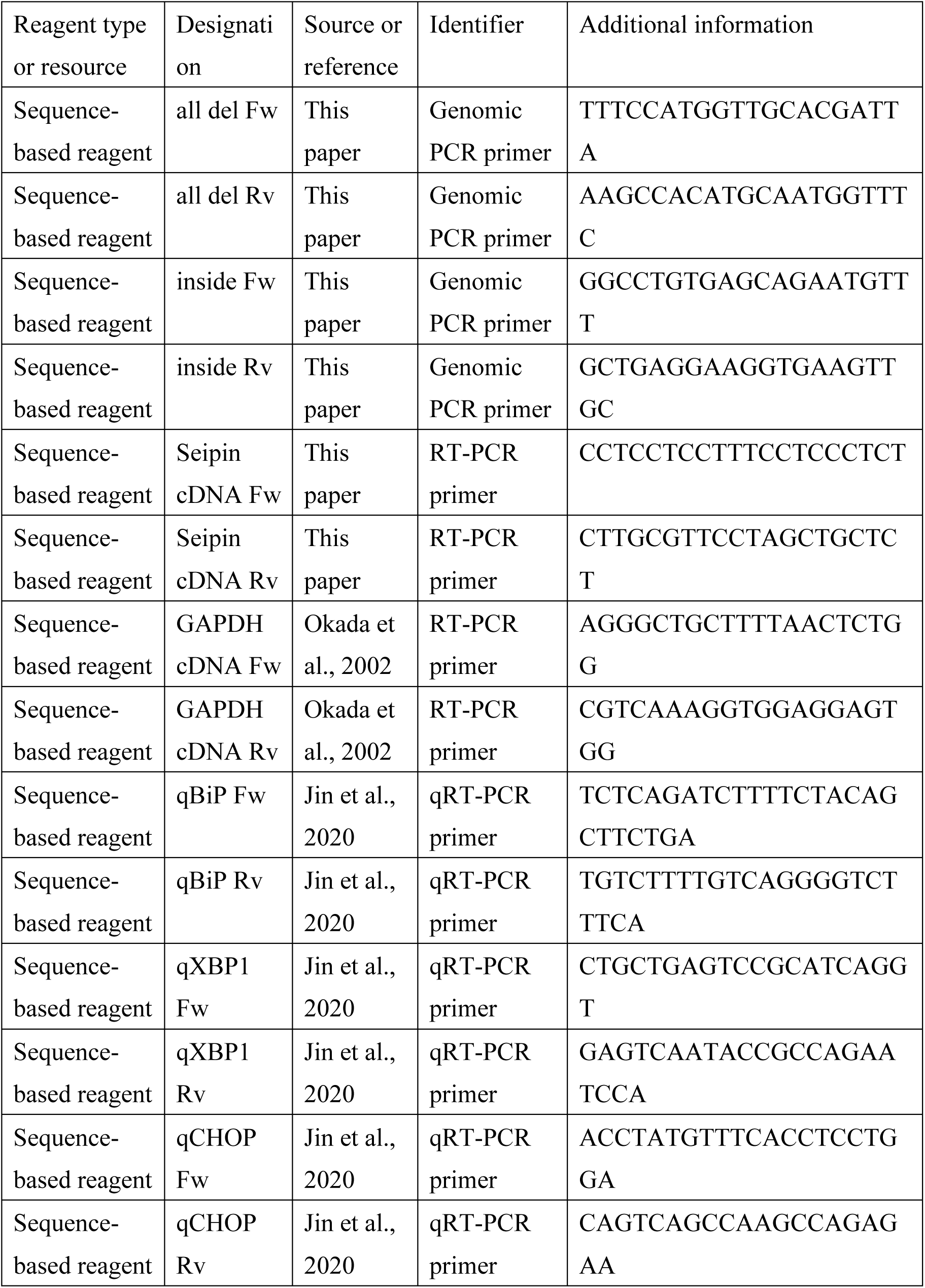
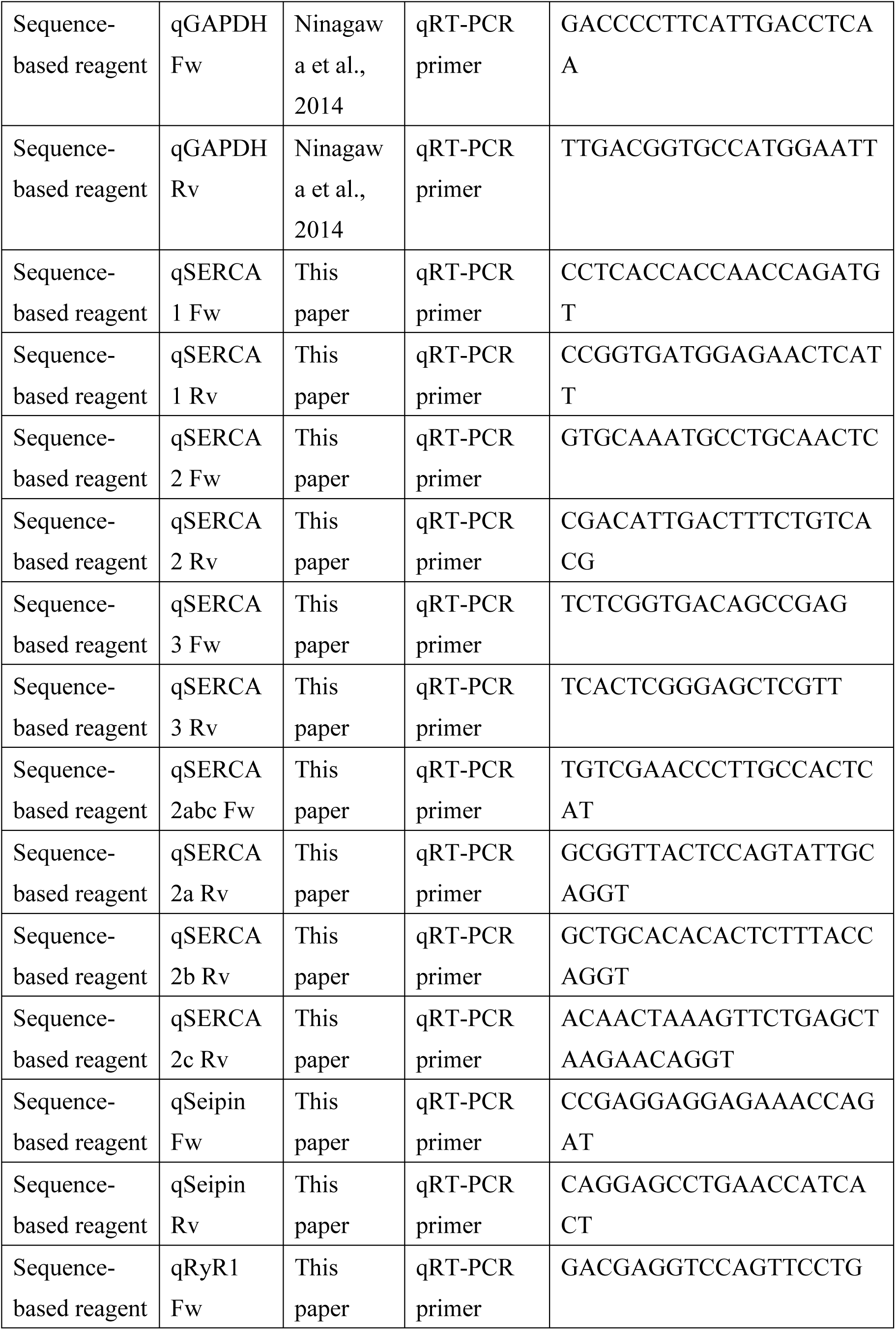
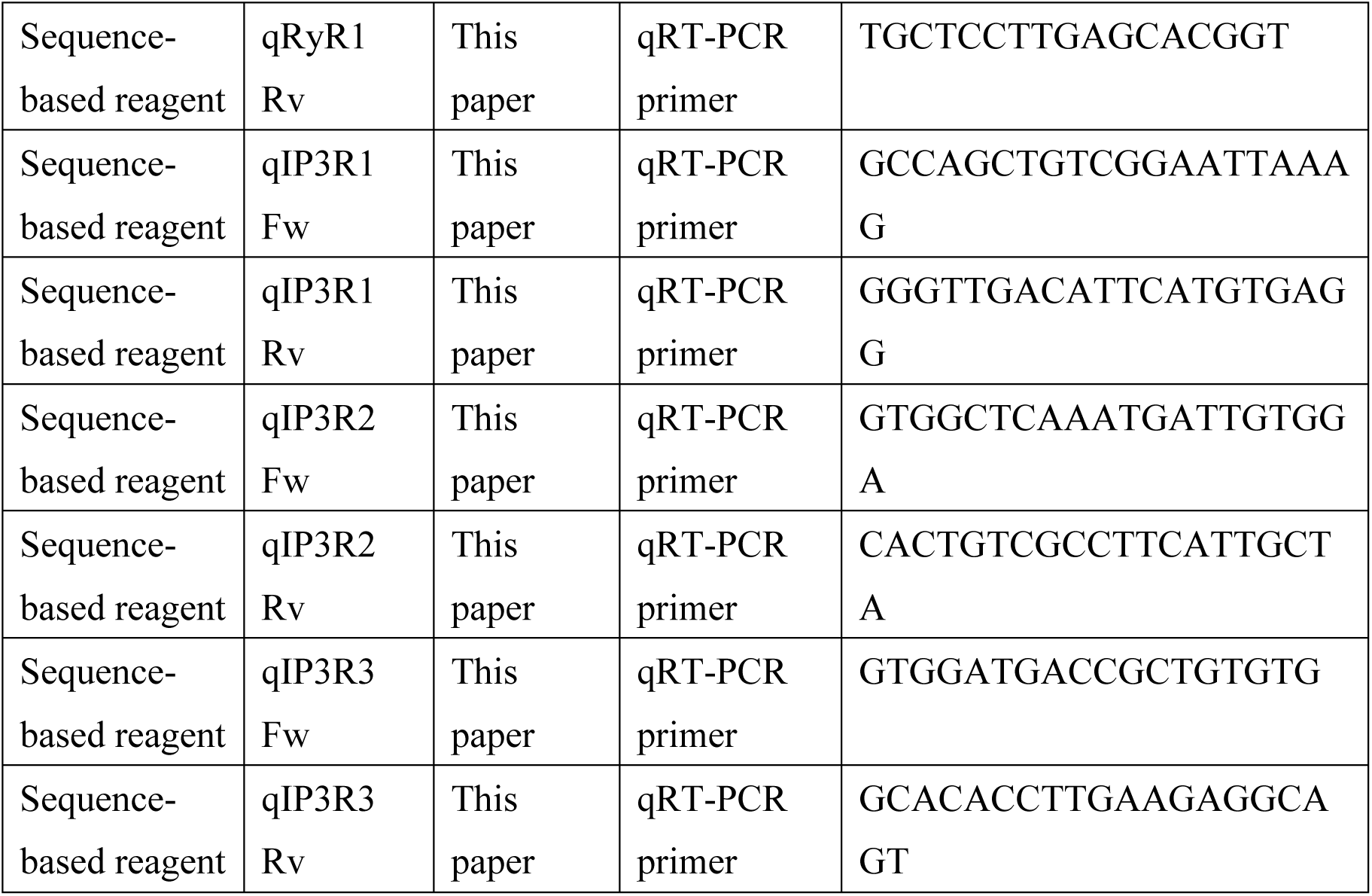

## ACKNOWLEDGEMENTS

The authors declare no competing financial interests. We thank Ms. Kaoru Miyagawa for her technical and secretarial assistance. This work was financially supported in part by grants from MEXT, Japan (19K06658 to T. I., 18K06216 to S. N., 17H06419 to K. M.) and AMED-CREST, Japan (20gm1410005 to K. M.)

---

**Fig. 1-figure supplement 1.**
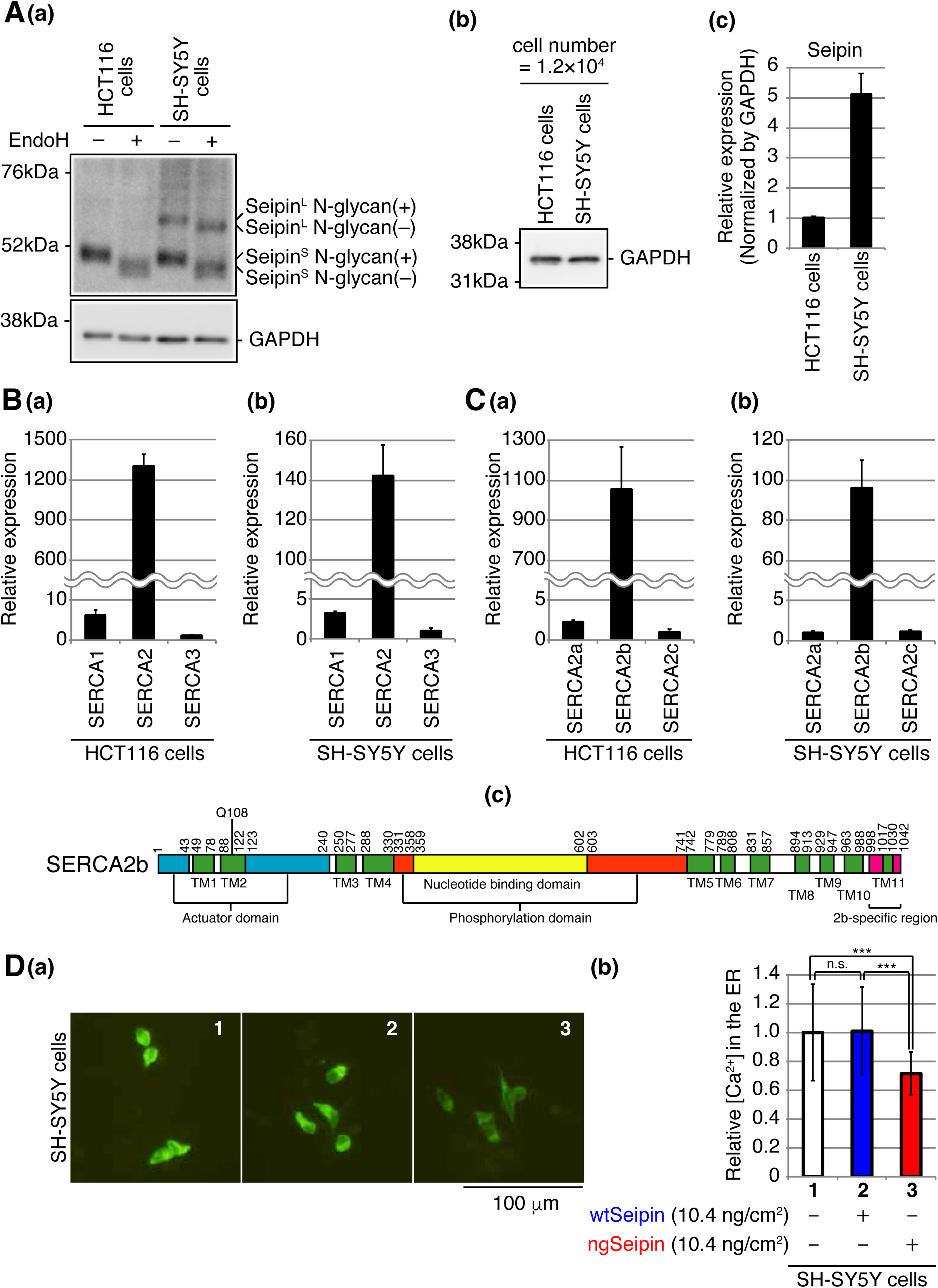
Effect of ngSeipin expression on calcium concentration in the ER of SH-SY5Y cells. (A) Lysates were prepared from HCT116 and SH-SY5Y cells, treated with (+) or without (-) EndoH, and analyzed by immunoblotting using anti-Seipin and anti-GAPDH antibodies. (b) Lysates were prepared from HCT116 cells and SH-SY5Y cells (12,000 cells each), and analyzed by immunoblotting using anti-GAPDH antibody. (c) Quantitative RT-PCR was conducted to determine the level of endogenous Seipin mRNA (a region shared by Seipin^L^ and Seipin^S^ was amplified) relative to that of GAPDH mRNA in HCT116 and SH-SY5Y cells (n=3). (B) Quantitative RT-PCR was conducted to determine the levels of endogenous mRNA encoding SERCA1, SERCA2 and SERCA3 relative to that of GAPDH mRNA in HCT116 (a) and SH-SY5Y (b) cells (n=3). The mean value of SERCA3 mRNA is set as 1. (C) Quantitative RT-PCR was conducted to determine the levels of endogenous mRNA encoding SERCA2a, SERCA2b and SERCA2c relative to that of GAPDH mRNA in HCT116 (a) and SH-SY5Y (b) cells (n=3). The mean value of SERCA2c mRNA and SERCA2a mRNA is set as 1 for (a) and (b), respectively. (c) Structure of human SERCA2b is schematically shown. Green, blue, orange, yellow and red boxes denote transmembrane (TM1-TM11) domains, actuator domain, phosphorylation domain, nucleotide binding domain, and SERCA2b-specific region, respectively. (D) (a) Fluorescent microscopic analysis of SH-SY5Y cells transfected with plasmid (104 ng/cm^2^) to express Cepia1er together with or without plasmid (10.4 ng/cm^2^) to express Myc-tagged wtSeipin or ngSeipin was conducted. Scale bar: 100 μm. (b) 15-17 pictures obtained from three independent experiments (5-7 pictures each) were quantified and are expressed as in Fig. 1E(b).

**Fig. 2-figure supplement 1.**
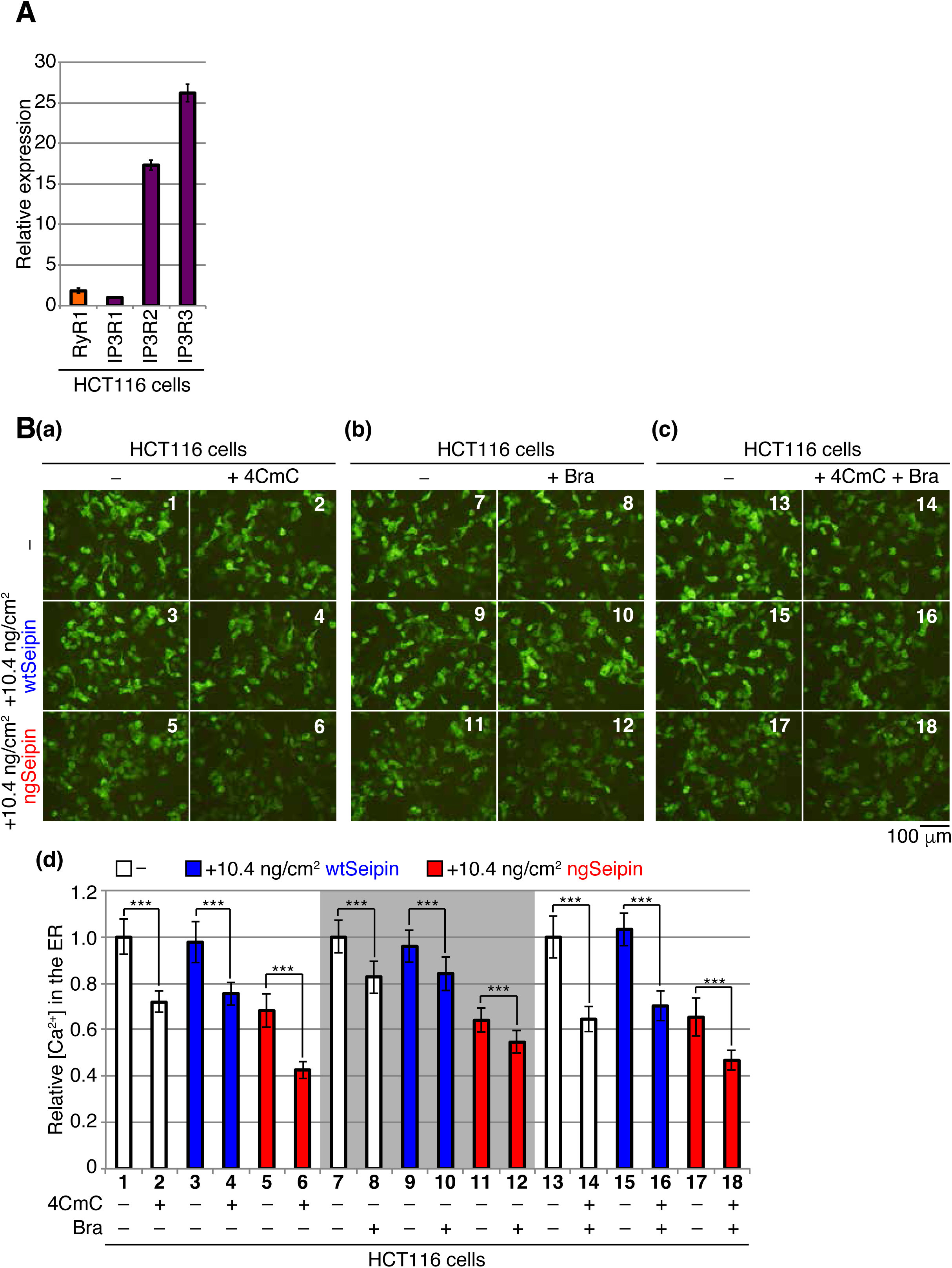
Effect of ngSeipin expression on RyR and IP3R. (A) Quantitative RT-PCR was conducted to determine the levels of endogenous mRNA encoding RyR1, IP3R1, IP3R2 and IP3R3 mRNA relative to that of GAPDH mRNA in HCT116 cells (n=3). The mean value of IP3R1 is set as 1. Note that the expression of RyR2 and RyR3 was not detected by our RT-PCR analysis. (B) HCT116 cells transfected with plasmid (104 ng/cm^2^) to express Cepia1er together with or without plasmid (10.4 ng/cm^2^) to express Myc-tagged wtSeipin or ngSeipin were untreated (-) or treated with (a) 1 mM 4-chloro-m-cresol (4CmC), (b) 10 μM bradykinin (Bra), or (c) both for 10 min. Fluorescent microscopic analysis was conducted. Scale bar: 100 μm. (d) Fluorescence intensities were quantified at each time point and are expressed as in Fig. 1E(b) (n=3). The experiments (a), (b) and (c) were conducted separately.

**Fig. 2-figure supplement 1.**
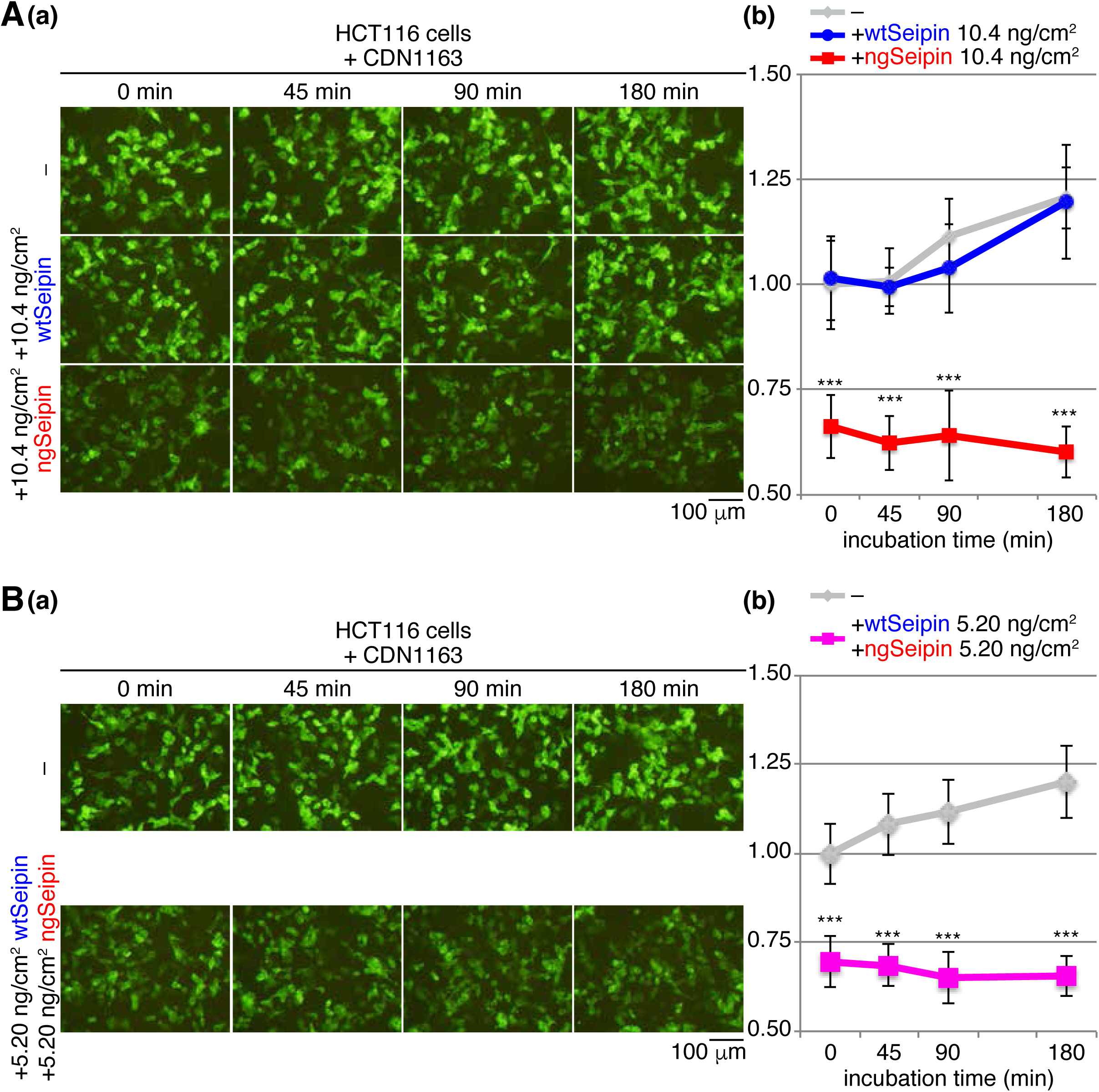
Effect of CDN163 on calcium concentration in the ER. (A) HCT116 cells transfected with plasmid (104 ng/cm^2^) to express Cepia1er together with or without plasmid (10.4 ng/cm^2^) to express Myc-tagged wtSeipin or ngSeipin were treated with 10 μM CDN1163 for the indicated times. (a) Fluorescent microscopic analysis was conducted. Scale bar: 100 μm. (b) Fluorescence intensities were quantified and are expressed as in Fig. 1E(b) (n=3). (B) HCT116 cells transfected with plasmid (104 ng/cm^2^) to express Cepia1er together with or without plasmid (5.20 ng/cm^2^) to express Myc-tagged wtSeipin plus plasmid (5.20 ng/cm^2^) to express Myc-tagged ngSeipin were treated with 10 μM CDN1163 for the indicated periods. (a) Fluorescent microscopic analysis was conducted. Scale bar: 100 μm. (b) Fluorescence intensities were quantified at each time point and are expressed as in Fig. 1E(b) (n=3).

**Fig. 3-figure supplement 1.**
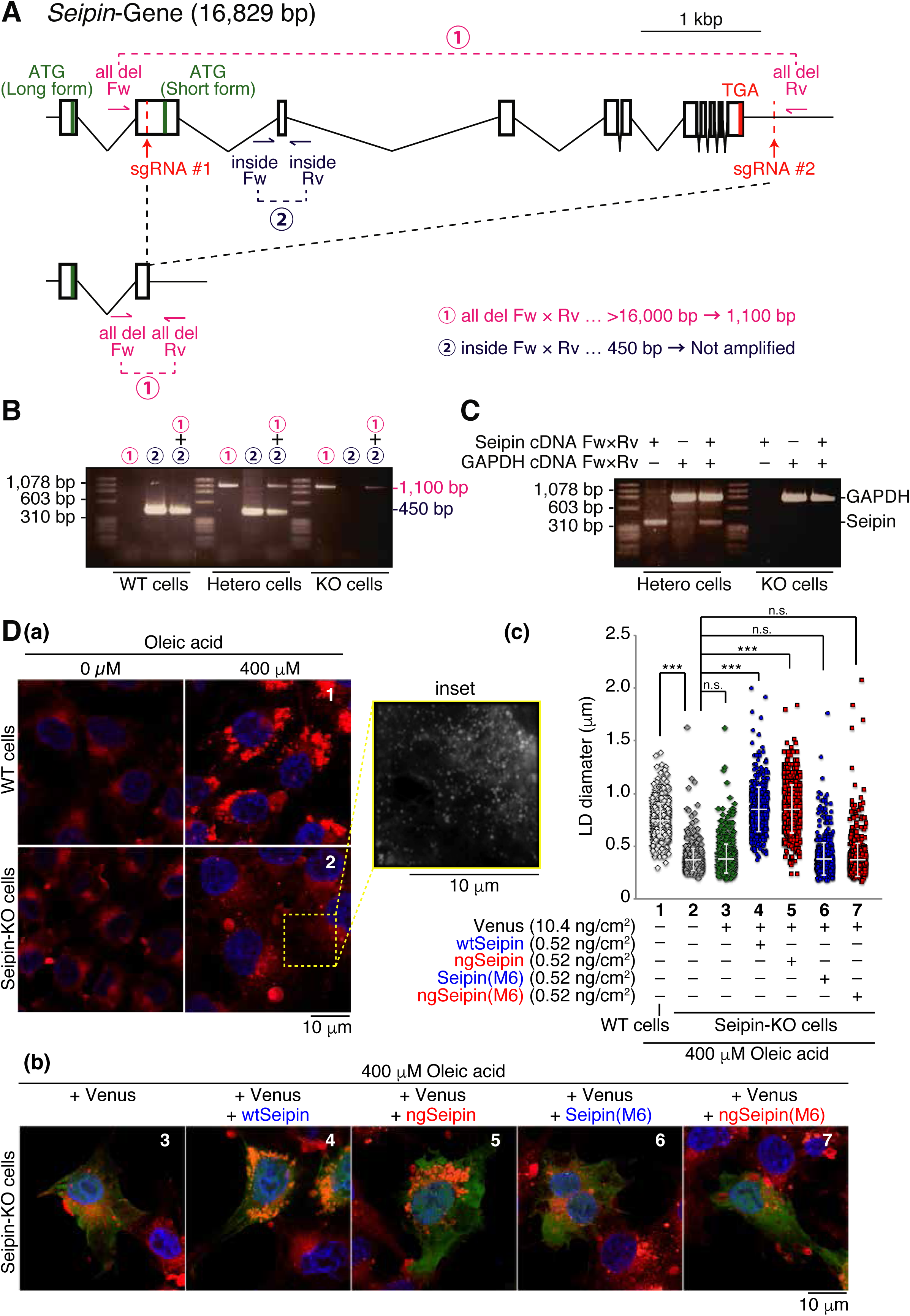
Construction and characterization of Seipin-KO cells. (A) Strategy of the CRISPR/Cas9-mediated cleavage to delete almost the entire the *Seipin* gene is shown. (B) Genomic PCR was carried out to confirm non-homologous end joining after cleavage. The positions of primers and expected sizes of amplified fragments are shown in (A). (C) Total RNA was prepared from heterozygous and Seipin-KO cells, and subjected to RT-PCR to amplify cDNA corresponding to Seipin and GAPDH mRNA. (D) (a) WT and Seipin-KO cells were untreated or treated with 400 μM oleic acid for 24 h, fixed and stained with Nile Red, and analyzed by fluorescent microscopy (Airyscan). Scale bar: 10 μm. (b) Seipin-KO cells transfected with plasmid to express Venus together with plasmid to express Myc-tagged wtSeipin, ngSeipin, Seipin(M6) or ngSeipin(M6) were treated with 400 μM oleic acid for 24 h and analyzed as in (a). (c) Diameters of total 551-908 lipid droplets [LD, see the inset in (a)] in 15-21 cells obtained from three independent experiments for (a) and (b) were measured and are shown.

**Fig. 6-figure supplement 1.**
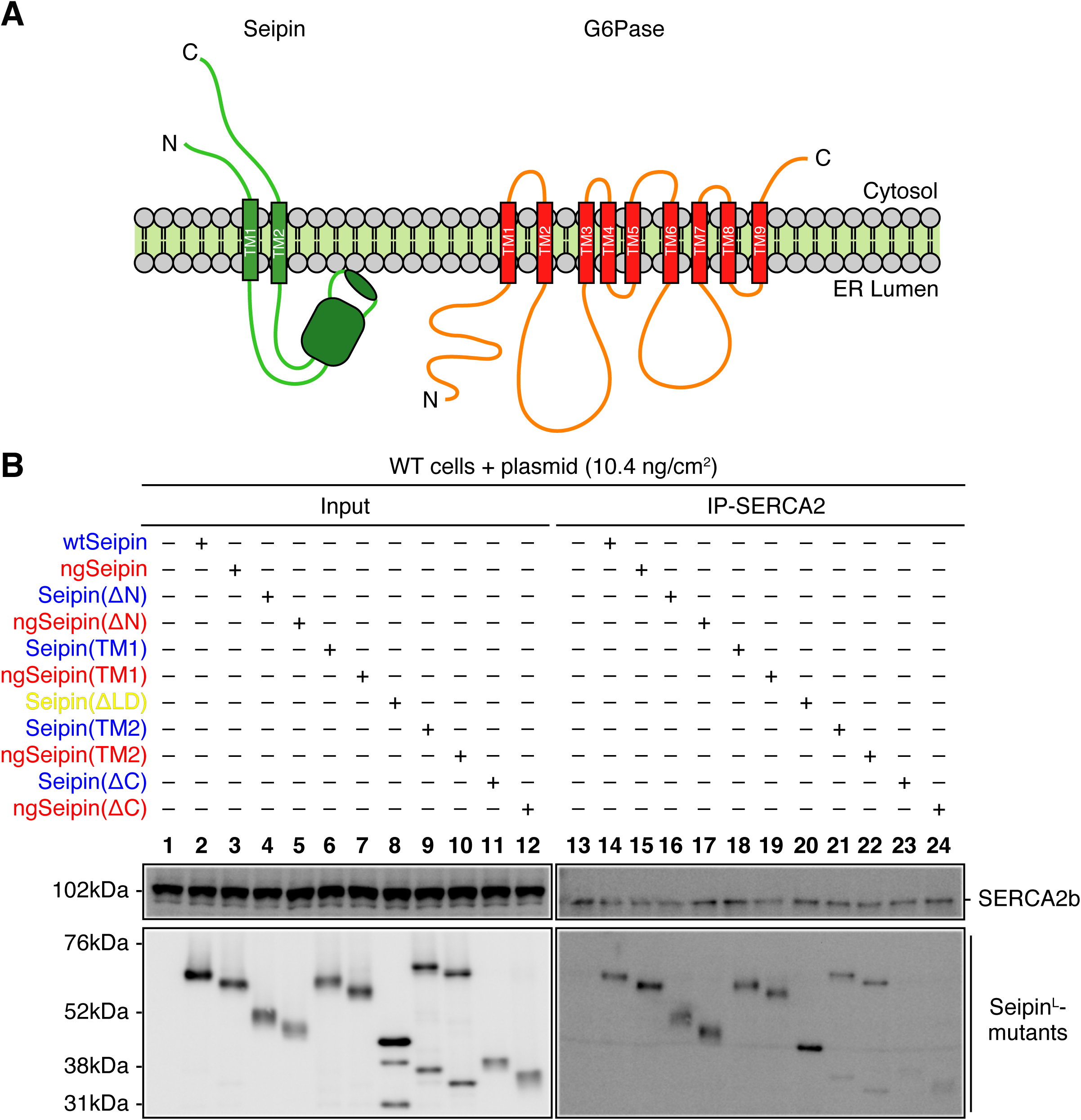
Effect of various deletions or replacements of ngSeipin on interaction with SERA2. (A) Structures of Seipin and glucose 6-phosphatase (G6Pase) are schematically shown. (B) Cell lysates were prepared from WT cells transfected with plasmid (10.4 ng/cm^2^) to express Myc-tagged wtSeipin or various mutant Seipin, subjected to immunoprecipitation using anti-SERCA2 antibody, and analyzed as in Fig. 1D.

